# Soil protists can actively redistribute beneficial bacteria along *Medicago truncatula* roots

**DOI:** 10.1101/2021.06.16.448774

**Authors:** Christopher J. Hawxhurst, Jamie L. Micciulla, Charles M. Bridges, Mikhael Shor, Daniel J. Gage, Leslie M. Shor

## Abstract

The rhizosphere is the region of soil directly influenced by plant roots. The microbial community in the rhizosphere includes fungi, protists, and bacteria, all of which play a significant role in plant health. The beneficial bacterium *Sinorhizobium meliloti* infects growing root hairs on nitrogen-starved leguminous plants. Infection leads to the formation of a root nodule, where *S. meliloti* converts atmospheric nitrogen to ammonia, a usable form of nitrogen for plants. *S. meliloti* is often found in biofilms and travels slowly along the roots, leaving developing root hairs at the growing root tips uninfected. Soil protists are an important component of the rhizosphere system who prey on soil bacteria and have been known to egest undigested phagosomes. We show that the soil protist, *Colpoda sp*., can transport *S. meliloti* down *Medicago truncatula* roots. By using pseudo-3D soil microcosms, we directly observed the presence of fluorescently labelled *S. meliloti* along *M. truncatula* roots and tracked the displacement of the fluorescence signal over time. Two weeks after co-inoculation, this signal was detected 52 mm, on average, farther down the roots when *Colpoda sp*. was also present compared with the experimental treatment that contained bacteria but not protists. Direct counts also showed that protists are required for viable bacteria to reach the deeper sections of root systems in our microcosms. Facilitating bacterial transport may be an important mechanism whereby soil protists promote plant health. As a sustainable agriculture biotechnology, protist-facilitated transport has the potential to boost efficacy of bacterial inoculants, thereby helping growers avoid overuse of nitrogen fertilizers and enhance performance of climate-smart, no-till farming practices.

**Importance:** Soil protists are an important part of the microbial community in the rhizosphere. Plants grown with protists fare better than plants grown without protists. Mechanisms through which protists support plant health include nutrient cycling, alteration of the bacterial community through selective feeding, and consumption of plant pathogens. Here we provide data in support of an additional mechanism: protists act as transport vehicles for bacteria in soil. We show that protist-facilitated transport can deliver plant-beneficial bacteria to the growing tips of roots that may otherwise be sparsely inhabited with bacteria originating from a seed-associated inoculum. By co-inoculating *Medicago truncatula* roots with both *S. meliloti*, a nitrogen fixing legume symbiont, and *Colpoda sp*., a ciliated protist, we show substantial and statistically significant transport with depth and breadth of bacteria-associated fluorescence as well as transport of viable bacteria. Co-inoculation with shelf-stable encysted soil protists may be employed as a sustainable agriculture biotechnology to better distribute beneficial bacteria and enhance the performance of inoculants.

## Introduction

The rhizosphere is the zone of soil surrounding plant roots that is under the influence of the root. This soil fraction contains 5 to 100 times more organisms per unit volume than adjacent bulk soil (1).Up to 40% of carbon assimilated by plants is contributed back to the soil as rhizo-deposits (2).In return for these carbon-rich root exudates, plants often benefit from microbial activity through a variety of mechanisms such as moisture retention, pathogen suppression, plant-hormone synthesis and the release of recalcitrant nutrients (3, 4). Furthermore, some soil dwelling bacteria, such as those from the family Rhizobiaceae, beneficially infect leguminous roots and directly provide fixed nitrogen in exchange for TCA cycle intermediates.

The functioning of soil and rhizosphere systems emerges, in part, from biological communities interacting in a spatially structured environment (5–12). In trying to better understand the importance of biological factors on the properties of soil, research has been dedicated to the study and interaction of different bacterial species and plants in spatially-constrained systems (6, 9, 10). Protists form an additional, important, and often overlooked component of the soil system. The presence of protists in addition to bacteria can promote and enhance size and weight of plant roots and shoots versus the presence of bacteria alone (13). Protist-associated enhancement to plant health can arise through several different mechanisms. In the “microbial loop” hypothesis, protists recycle limiting nutrients by grazing on the microbes fed by root exudates (14–18). Protists may also alter the microbial composition of the rhizosphere through selective grazing (18, 19). A third potential mechanism, investigated here, is that protists may improve plant health by acting as a transport facilitator: directly relocating bacteria or other “cargo” along downward-growing roots (20–22). Similar examples include transport of fungal spores hitchhiking on bacterial flagella (23). Other examples are given in recent reviews (24, 25).

Plant growth-promoting rhizobacteria (PGPR) are beneficial soil bacteria used in agriculture to improve crop yield and disease resistance. PGPR-mediated nutritional effects include symbiotic nitrogen fixation, non-symbiotic nitrogen fixation, release of recalcitrant nitrogen from soils, phosphate solubilization, and iron delivery (26–29). Commercially-used PGPR include many species of nitrogen-fixing bacteria: *Azospirillum* spp., *Azotobacter* spp., *Pseudomonas* spp., *Bacillus subtilis* strains, *Bacillus megaterium* strains and *Trichoderma* spp. (a fungal genus) (29). Many additional species are under investigation in laboratory, greenhouse, and field trials, but have not yet been commercialized.

PGPR are typically inoculated onto seeds before planting or they are directly applied to the soil during planting. Soil bacteria usually grow as biofilms and spread very slowly. Thus, bacteria inoculated onto, or near, seeds often do not leave the area of inoculation and do not establish a population on the root large enough to provide beneficial services (30–33). Further, symbiotic infection of root hairs with nitrogen-fixing rhizobia, such as *S. meliloti*, often occurs at root hairs located near root tips. Because roots grow only near their tips, suitable locations for infection in a seedling are located further and further from the soil surface as a seedling develops. A lack of bacteria at the root tip will most often result in decreased rate of symbiotic infection and nodulation (34).

Soil protists have the potential to act as transport vehicles for bacteria. Soybean roots can grow at ∼3000 µm per hour (35). For comparison, the spreading rate of biocontrol *P. fluorescens* biofilms is less than 10 µm per hour (36), while ciliated protists can swim at speeds up to 400 µm per second for short periods (37). We have previously shown that protists move fluorescently-labeled microspheres and bacteria in emulated soil micromodels (20). In situ, protists are known to carry bacteria, most commonly as intracellular parasites, for example when *Legionella* spp. infect aquatic amoeba or *Campylobacter* spp. infect ciliates (38, 39). Also, bacterial transport by protists has been demonstrated in the soil amoeba *Dictyostelium discoideum*, which can farm bacteria and carry them to new habitats (40, 41). Although protists graze on bacteria, ingestion is not always fatal (38). Further, bacteria have been observed attached to the outer surfaces of protists, thus providing a means for transport that does not require ingestion (20). We propose protists contribute to the rhizosphere community, in part, through their capacity to move bacteria. The goal of this study was to test the hypothesis that soil protists, despite grazing, can enhance dispersal of bacteria along root systems. We measured the spatial distribution of fluorescently labelled *S. meliloti* through soil-containing mesocosms over time in the presence and absence of a soil-isolated, ciliated, protist from the genus *Colpoda*. Our methods included both nondestructive direct imaging repeated over time and direct counts of viable bacteria and protists collected from root systems at the conclusion of each experiment.

## Experimental Materials and Methods

### Model soil

Sterile model soil was comprised of play sand (Quikrete) and fine vermiculite combined 1:1 in a soil mixer. The mix was fractionated by sieving and the 125-250 µm fraction and 63-125 µm fraction were combined 1:2 to emulate the particle sized distribution of a model sandy loam (11).

### Plants

*Medicago truncatula* is a model legume that offers uniform growth and natural symbiosis with *S. meliloti*. To synchronize the plants, *M. truncatula* cv. Jemalong seeds were sterilized by successive washing with 95% ethanol for 30 min, 10% bleach for 15 min, and sterile diH_2_O for 15 min on a shaking platform at 80 rpm. Next, seeds were imbibed in sterile diH_2_O in the dark for 24 h. Finally, seeds were rinsed in sterile diH_2_O, transferred to an empty, sterile plastic Petri dish, and held inverted in darkness overnight for germination.

### Bacterial strain construction

The BioFAB plasmid pFAB2298 was kindly provided by Vivek Mutalik (42). We amplified the expression cassette containing Ptrc-BCD2-mRFP1 with Phusion polymerase (Fisher) as a single unit from pFAB2298 using primers 5’-ATCTGCAGGCTTCCCAACCTTACCAG-3’ and 5’-CAGTATCGATAGTCAGTGAGCGAGGAAGC-3’, which contain 5’ *Pst*I and *Cla*I restriction sites, respectively. We digested the resulting 1.244 kb amplicon with *Pst*I and *Cla*I (New England Biolabs) and cloned it into pBlueScript II SK(+) to create pCMB21. We digested the expression cassette from pCMB21 using AvrII and EcoRI-HF (New England Biolabs) and cloned into pSEVA551 (*tetA*, RSF1010 *oriV*) to create pCMB35 (43). The plasmid pCMB35 contains the broad host range RSF1010 origin of replication, the *tetA* gene encoding a tetracycline efflux transporter which confers tetracycline resistance, and a protein expression cassette which employs the strong, constitutive Ptrc promoter and a bicistronic 5’ UTR region with two ribosome binding sites (RBS) which allows even expression levels across expressed genes, in this case the gene *mRFP1*. Constitutive mRFP1 expression by Rm1021 allows the location of bacteria within the device to be identified through fluorescence microscopy. We introduced plasmid pCMB35 into *S. meliloti* strain Rm1021 by electroporation in a 1 mm gapped cuvette at 2200 V and plated on TY medium (6 g/L tryptone (BD), 3 g/L yeast extract (BD), 0.38 g/L CaCl_2_ (Acros), 15 g/L agar (BD)), amended with 500 μg/mL streptomycin sulfate (Acros) and 5 μg/mL tetracycline hydrochloride (Sigma).

### µ-rhizoslide construction

We constructed small-scale, soil-containing, mesocosms adapted from the rhizoslide designed by Le Marié, *et al* (44). Mesocosms were comprised of a 3D printed spacer with a hollow opening which was filled with model soil. We refer to our version as a “µ-rhizoslide.” After several design iterations to balance space for root growth, plant health, and imaging consistency, the final spacer design was 0.5 cm thick, featuring a 1.5 cm wide, 7.5 cm long main channel with a 1.5 mm connector at the top of the channel to stiffen the spacer and limit deformation (Figure 1).

**Figure 1.**
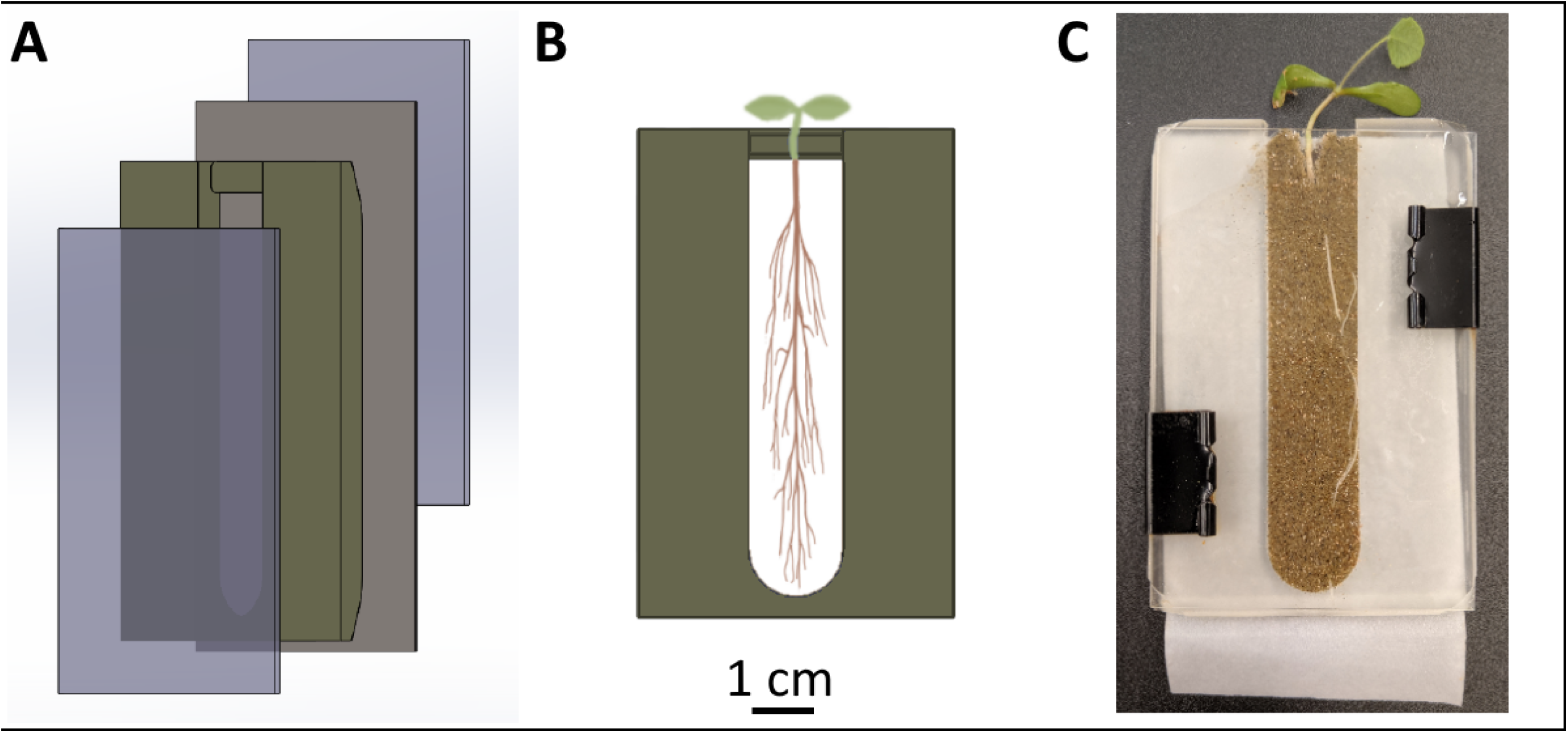
µ-rhizoslide construction and use. A) Computer-aided design (CAD) drawing of the rhizoslide assembly showing layers including 3D printed rhizoslide spacer and filter paper wick between two glass slides. B) Schematic of assembled µ-rhizoslide. C) Photograph of µ-rhizoslide in use.

Our µ-rhizoslides facilitate direct optical imaging of the growing plant root system while minimizing water-based transport effects. The µ-rhizoslide assembly consists of a 3D printed spacer sandwiched between two 50 × 75 mm glass slides, with a 50 × 85 mm filter paper wick located between the back of the 3D printed spacer and the rear microscope slide to allow for plant hydration via capillary action without gravity driven, bulk water flow. Spacers were designed using Solidworks 2018, imported into PreForm software and printed on a Formlabs Form 2 stereolithography 3D printer using Clear Resin GPCL04 (Formlabs, Somerville, MA).

To improve biocompatibility, printed spacers were treated to leach labile resin or crosslinker using a protocol adapted from Kadilak *et al*. (45). Treatment is necessary because un-crosslinked monomer can have antimicrobial properties (46). Briefly, we soaked 3D printed spacers in an isopropanol bath for 30 min, then rinsed with additional isopropanol from a squeeze bottle. This procedure was repeated twice more to remove un-crosslinked resin before resting in a 60 °C oven for 60 min.

Assembly of µ-rhizoslides proceeded as follows. The 3D printed spacer resting on a glass slide was filled with sterile diH_2_O-saturated sterile model soil, then covered with a filter paper wick, and finally the µ-rhizoslide assembly was closed by affixing a second glass slide to the back. Germinated seeds were placed in a small depression at the top of the soil column. We kept the µ-rhizoslides separated by treatment in transparent plastic boxes with each µ-rhizoslide positioned 45° off vertical with the filter paper wick above the soil to encourage root growth directly against the glass on the front of the assembly. This arrangement also helped protect the roots from light exposure, as we kept the µ-rhizoslide boxes in a growth chamber under a 16:8 h light:dark regime at 23 °C. We allowed the plants to grow in the µ-rhizoslides for 2 d to ensure plants became established prior to microbial inoculation and imaging.

### Growth of protists and bacteria

The protists used here were identified by 18s rRNA sequencing as a belonging to ciliate genus, *Colpoda*, which we refer to as UC1 (47). We isolated UC1 from a bean rhizosphere. UC1 was maintained in 25 cm^3^ tissue-culture flasks (Thermo Scientific #130192) with only 5 mL of Page’s saline solution (48) to promote oxygenation. For experimental runs, protists were fed heat-killed *E. coli* DH5a, inoculated at OD_595_ = 0.005 as measured in 100 µL of a suspension in a 96-well microtiter dish. Prior to plant inoculation, UC1 cultures were stored in darkness, at room temperature, for 3 weeks. Cysts were removed from the cell culture flask using a sterile cell scraper, washed twice with Page’s saline solution, then resuspended in 100 µL of the same solution. Protist cysts were washed, concentrated, and counted the day of inoculation. We estimated the concentration of the cysts by averaging the number of cysts counted in three 1 µL spots under a microscope. For experimental runs using trophozoites (protists in their active growing stage) encysted protist cultures were washed two days before inoculation to allow for excystment. Once most protists had emerged from the cysts and were active, they were concentrated and counted to enable delivery of a consistent number of protists per replicate.

Bacteria inoculants were prepared as follows. A single colony of *S. meliloti* strain Rm1021/pCMB35 was picked from a TY + tetracycline (5 µg/ml) plate, grown in a 125 mL flask containing 25 mL of liquid TY + tetracycline (5 µg/ml) at 30 °C with shaking at 160 rpm for 3 d. Cells were washed twice with Page’s saline solution then resuspended in Page’s to OD_595_ = 0.100. OD_595_ was determined in a 96-well microtiter dish using 100 µL of cell suspension.

### Experimental design and setup

For each of the three experimental runs, 2-day seedlings were inoculated directly at the shoot-root interface with 10 µL of one of three different solutions:

1. Page’s saline solution (Control);
2. Page’s saline solution containing 10^7^ CFU Rm1021/pCMB35 (Bacteria Only);
3. Page’s saline solution containing 10^7^ CFU Rm1021/pCMB35 + 1.5 × 10^3^ *Colpoda* UC1 (Bacteria + Protists).

Each treatment of each experimental run was replicated between 3 and 5 times for a total of 41 different µ-rhizoslides prepared and analyzed (see Table 1). To inoculate µ-rhizoslides, we positioned chambers at an angle either wick side up, the same way they were incubated, (Experimental Run 1) or wick side down (Experimental Runs 2 & 3) and held inverted for 1 hr. Inoculating wick down prevented the formation of a large fluorescence bloom directly against the forward-facing glass at the initial time point. Eliminating this large source of “background” fluorescence at the initial time point made it easier to measure subsequent changes in the fluorescence signal over time. Protists were either encysted (Experimental Run 1) or trophozoites (Experimental Runs 2 & 3); the use of trophozoites prompted quicker onset of protist-facilitated transport activity. The watering regime was either to water µ-rhizoslides weekly with complete fertilizer (Experimental Run 1) or every 4^th^ day, alternating between sterile diH_2_O and a nitrogen-free fertilizer (Experimental Runs 2 & 3). More frequent watering and reduced nutrient amendments helped plants avoid water starvation and fungal overgrowth. All replicates for all treatments were handled identically for a given experimental run and the effect of methods on results are discussed. In all cases, µ-rhizoslides were positioned during plant growth at a 45° angle with the filter paper above the soil (except when briefly repositioned for imaging, when the front face was positioned down on the stage of an inverted microscope). Experimental Run 1 lasted two weeks post inoculation, then was halted because several of the plants had died, prompting the modifications to the methodology described here. Experimental Runs 2 and 3 lasted three weeks post inoculation then were stopped because the roots in several replicates had reached the end of the soil-filled channel.

**Table 1.**
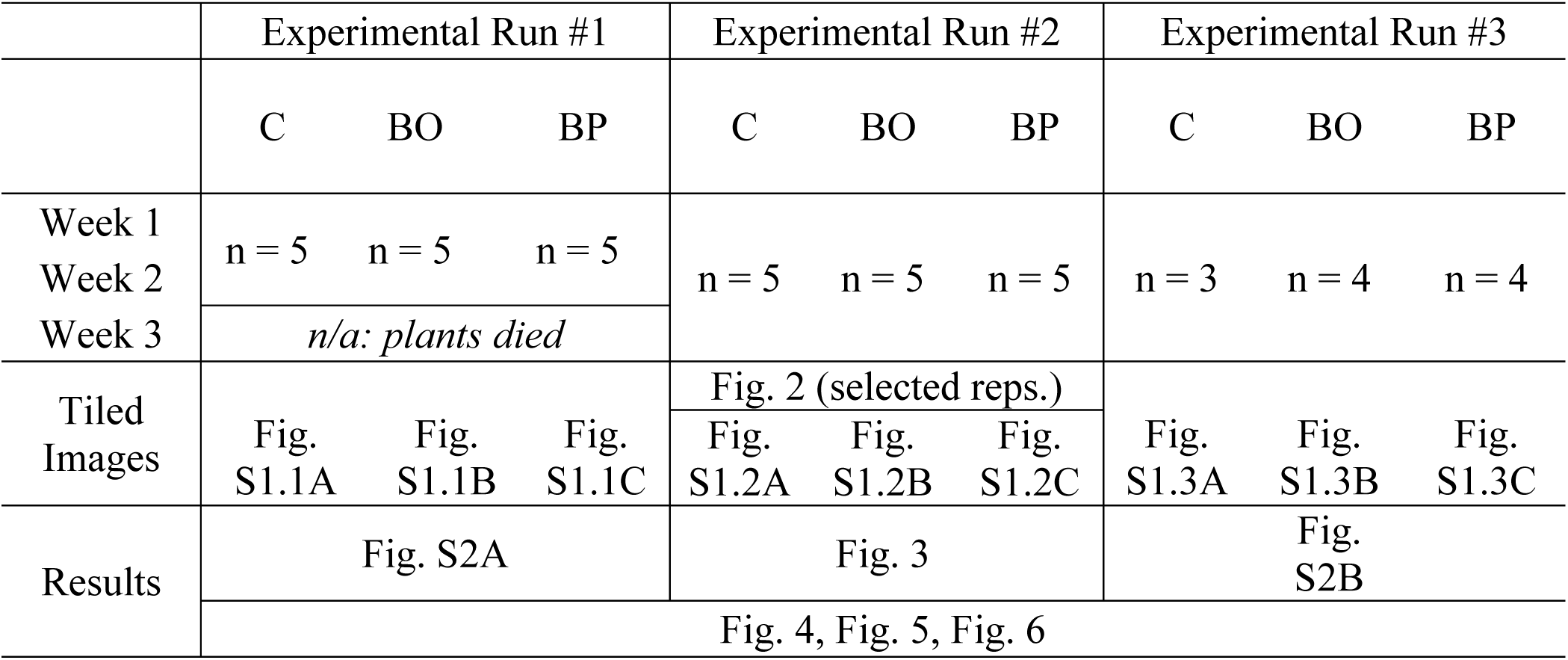
Experimental design showing three independent experimental runs each with multiple replicates of each treatment. A grand total of 41 separate µRhizoslides were imaged on successive weeks, then sacrificed to enumerate bacteria and protists. Treatment abbreviations are Control (C); Bacteria Only (BO); and Bacteria + Protists (BP). Plants in Experimental Run 1 died during the third week leading to modified conditions for Experimental Runs 2 and 3.

### Imaging and Image Analysis

On the day of inoculation and at intervals of 1 week subsequently we imaged the entire soil area of each µ-rhizoslide using a Zeiss AXIO-Observer Z1 inverted microscope (Carl Zeiss Inc., Germany). Collected images included tiled brightfield and fluorescence images using a 5× Zeiss EC Plan-Neofluar 5×/0.16 M27 lens and an AxioCam MRm Rev.3 camera (1348 × 1040 pixels, 6.45 × 6.45 µm/pixel) set to collect 585-binned photos (2 × 2 binning) when imaging. The transmission-based brightfield light equipped on the microscope could not illuminate through the 0.5 cm of soil, so we moved the brightfield lamp to an external stand and illuminated directly from the front of the µ-rhizoslide. For fluorescence imaging, we found 570 nm excitation and 620 nm emission filters gave the strongest fluorescence signal for *S. meliloti* Rm1021/pCMB35. We used the Zeiss Zen Pro 2.3 software to stitch a single 207-megapixel image (29,341 × 7,072 pixels) from each set of 585 photos then exported the tiled image as a 12-bit grayscale TIFF at full resolution.

We created composite images for each replicate of each experimental run to illustrate the movement of bacteria with and without protists (Figure 2, Figure S1). Here, brightfield images showing the fully-developed root were overlain with false-colored fluorescence images at each timepoint to show the progression of the fluorescence signal over time. These composite images are for illustration purposes only and were not used directly for quantitative image analysis.

**Figure 2.**
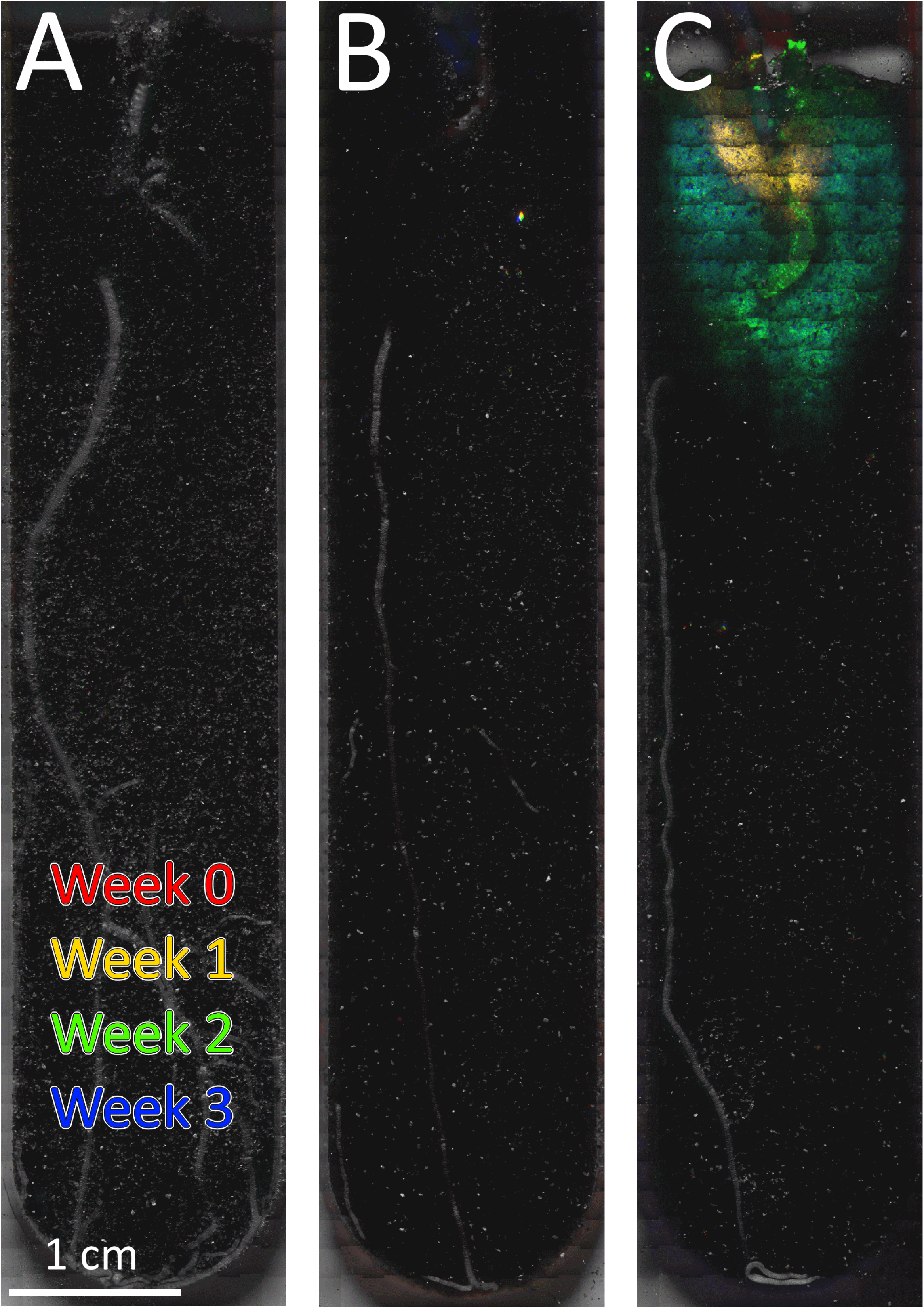
Selected composite brightfield (at last imaging date) and fluorescence (false-colored by week as indicated) images of soil channels for: A) Control: soil and plant inoculated with 10 µL buffer: no microbes added; B) Bacteria Only: soil and plant inoculated with 10 µL buffer containing 10^7^ CFU bacteria; C) Bacteria + Protists: soil and plant inoculated with 10 µL buffer containing 10^7^ CFU bacteria plus 1.5 × 10^3^ *Colpoda sp*. protists. Images for all experimental runs and all replicates are available in Figure S1.

For quantitative image analysis, stitched fluorescence images were imported into MATLAB and converted into 29,341 × 7,072 matrices. The initial (i.e., Week 0) tiled fluorescence image of each µ-rhizoslide was subtracted from the corresponding tiled fluorescence images from subsequent time points to remove background fluorescence, such as from mica particles in the soil. Any resulting negative pixel value was set equal to 0. Tiled images were cropped to remove the non-soil portion, eliminating fluorescence from the 3D printed spacers. As *S. meliloti* bacteria was predominately found along roots and towards the center of the soil channel, cropping out the region adjacent to the spacer had little effect on the bacterial fluorescence signal.

To quantify displacement of the fluorescence signal by treatment and time, we partitioned the photo matrixes into sections each approximately 1 mm high in the longitudinal direction (i.e., along the root axis) and extending across the full breadth of the soil-filled channel. The dimensions of each of these sections is 386 × 7,072 pixels. Next, we determined the proportion of each section with a non-zero fluorescence signal. We define “fluorescent pixel fraction” as the ratio of nonzero pixels to total pixels in a section.

We defined “longitudinal extent” as the depth of the 1-mm section furthest from the soil surface where fluorescent pixel fraction > 0.1 (i.e., where more than 10% of pixels in the section are non-zero, after correcting for background as described above). We defined “lateral extent” as the greatest fluorescent pixel fraction observed for a given replicate at a given point in time, regardless of depth, then multiplied this ratio by 1.5 cm to represent the width of a continuous fluorescent region.

The statistical significance of adding protists on both longitudinal and lateral extent was determined by nine binary comparisons for each (one for each pair of treatments in each week). Here, data from the three experimental runs are pooled together and p-values are computed using unpaired t-tests, two-sided, with unequal variance, and applying a Bonferroni correction for multiple hypothesis tests. The effect of experimental run on longitudinal and lateral extent was also determined using linear regression. This is important because of the small changes to the experimental procedures employed. For each direction and week, we regressed longitudinal extent or lateral extent on treatment and experimental run, with the control treatment and Experimental Run 1 as the omitted reference category. All statistical analyses were done using Stata SE 17 (StataCorp 2021, College Station, TX).

### Enumeration of *Colpoda sp*. and *S. meliloti* in µ-rhizoslide

After final imaging, µ-rhizoslides were disassembled and roots were separated from the bulk soil. Root systems (plant tissue plus adhering soil) were bisected into upper (“shallow roots”) and lower (“deeper roots”) segments beginning 2.5 cm below the root/shoot interface. Root weight was recorded, then each root segment was suspended in 10 × Page’s saline solution by weight in 50 mL conical tubes then vortexed for 1 min at medium speed on a tabletop vortex mixer.

For bacterial abundance, 10-fold serial dilutions of each vortexed root extract were prepared in Page’s saline solution then 10 µL spots of each dilution were placed onto TY + tetracycline (5 µg/mL) agar plates in square grid-marked Petri dishes and incubated at 30 °C for 3 d. Colonies were counted, and bacterial abundance of *S. meliloti* strain Rm1021/pCMB35 on each root system segments was estimated by multiplying colony counts by the corresponding dilution factor.

To estimate protist abundance, 3-fold serial dilutions were carried out with octuple replication in 96-well microtiter dishes using Page’s saline solution + heat-killed *E. coli* DH5a (OD_595_ = 0.050). Microtiter dishes were incubated at room temperature in darkness for 1 week.

The presence or absence of protists in each well was recorded. For analysis, we assumed “at least one” protist made it into any well that contained a population of protists after one week. The Minimum Recovered Number per ml (MRN/ml) of protists was calculated as follows: MRN/ml = 10 * 3^(farthest positive dilution factor – 1)^. Significance was determined as described previously with paired or unpaired t-tests as noted in the data tables.

## Results and Discussion

### Performance of µ-rhizoslides

The µ-rhizoslides described were constructed as a smaller version of the rhizoslides designed by C. L. Marie (44) but optimized for microscopy and increased throughput. We report root growth and plant-microbe interactions were readily monitored on a standard inverted microscope using these smaller µ-rhizoslides. *M. truncatula* roots occupied a region 15 × 75 mm and were imaged as a series of overlapping tiled images. Brightfield microscopy was used to localize root tissue including visualization of primary, secondary and tertiary roots, and fluorescence microscopy was effective in monitoring displacement of a bacteria-associated fluorescence signal over time. Images were collected once per week for three weeks after planting, and in our case, experimental duration was limited only by growth of the plant. The µ-rhizoslide system described here is simple and inexpensive to produce, can support seeds as large as 4 mm diameter, and can support experiments until root growth extends beyond the chamber size.

### Displacement of *S. meliloti* is enhanced by protists

In order to test the hypothesis that addition of protists can enhance dispersal of bacteria along root systems, the nitrogen fixing symbiont *S. meliloti* strain Rm1021 was transformed with plasmid pCMB35 which constitutively expressed mRFP. The movement and growth of Rm1021/pCMB35 was nondestructively assayed by fluorescence microscopy, and the role of protists in facilitating its transport was determined by comparing displacement of the fluorescence signal over time across treatments. Typical of our results (Figure 2) we observed fluorescence gradually spreading with depth and breadth along and across the growing roots. Strikingly and repeatedly, we observed that the fluorescence signal reached further down the primary axis of the growing roots (i.e., in the longitudinal direction) and spread more side-to-side when in the presence of soil protists. Tiled images for all forty-one µ-rhizoslides at all points in time are available in the Supplemental Materials, Figure S1.

To analyze these images quantitatively, fluorescent pixel fraction (ratio of non-zero to total pixels) was quantified for thin vertical strips of the soil channel for each composite image. A plot of fluorescent pixel fraction versus depth in the chamber illustrates the displacement of the fluorescence signal in the longitudinal direction over time (Figure 3 and Figure S2). The results show small values of fluorescent pixel fraction scattered fairly uniformly with depth for Control and Bacteria Only treatments, with much larger values for fluorescent pixel fraction in all replicates of the Bacteria + Protists treatments, but in particular in the shallow portion of the soil channel, and at later points in time. For example, for Experimental Run 2 at Week 2, most replicates show a wide band spanning 20-30 mm of depth where fluorescent pixel fraction > 0.5. At Week 2 and Week 3 some replicates show appreciable fluorescent pixel fraction values extending all the way to the end of the soil-filled channel. Meanwhile, for the other two treatments (Control, where no microbes are added, and Bacteria Only), no replicate showed such marked fluorescence.

**Figure 3.**
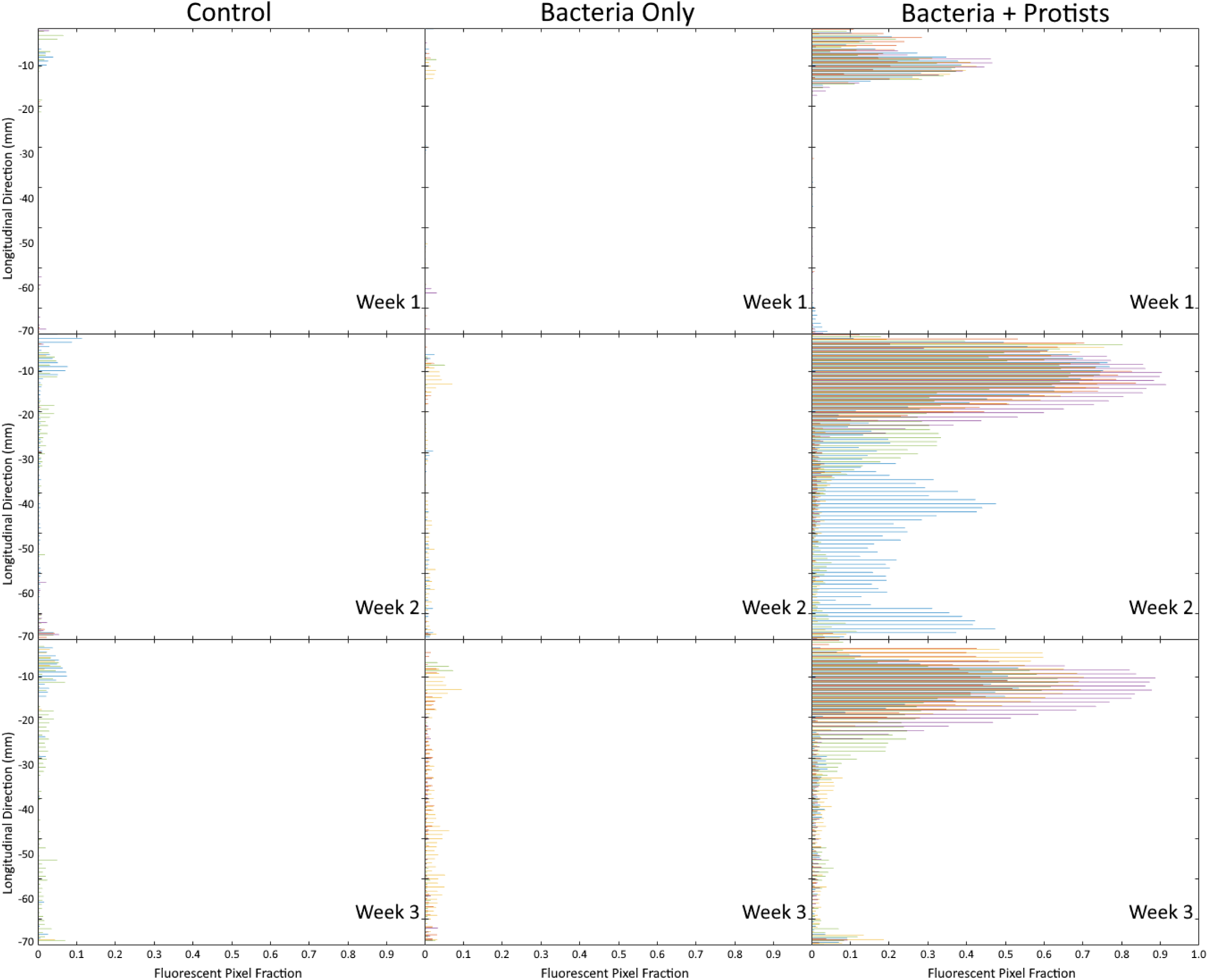
Spatial distribution of fluorescence signal over time. For each treatment (arranged above in columns), the entire soil channel was imaged at different points in time (Week 0, Week 1, Week 2, Week 3). Here, “Fluorescent Pixel Fraction” (the ratio of non-zero pixels to total pixels) was computed for binned strips (each measuring 1 mm high and 15 mm wide) of the soil channel after background was subtracted (Week 0) and plotted with distance in the longitudinal direction. Data from Experimental Run 2 with replicates distinguished by color. Full data from the other two experimental runs can be found in Supplemental Material, Figure S2.

The pattern of fluorescent pixel fraction versus depth (i.e., the longitudinal direction) in Experimental Run 3 is quite similar to Experimental Run 2, with the greatest fluorescent pixel fraction near the surface. For Experimental Run 1 the fluorescence pattern is somewhat different, with much lower fluorescent pixel fraction values overall, yet still substantially higher values of fluorescent pixel fraction in the Bacteria + Protists treatment than in the Control or the Bacteria Only treatments. Referencing back to the images Experimental Run 1, Bacteria + Protists Treatment (Figure S1.1C), shows in all five replicates marked fluorescence directly adjacent to root tissue and extending along the full depth of the channel: this would result in low fluorescent pixel fraction values even through the root may be fully colonized.

Because even a small number of bacteria that reach a favorable habitat can create robust populations in time, the spatial extent of even a relatively small fluorescence signal is perhaps more important to our analysis than the total amount of fluorescence. We defined “longitudinal extent” as the greatest depth where fluorescent pixel fraction > 10%. Practically, exceeding this threshold of 10% fluorescent pixels reduces the probability of spurious results and conceptually may indicate some potential for bacterial colonization. A plot of longitudinal extent by replicate and time (Figure 4) show the effect of protists on mobilizing bacteria across the entire root system. For example, by Week 2 the median longitudinal extent in the Bacteria + Protists treatment was 52 mm further down the root as compared with the Bacteria Only treatment. By Week 3, while replicates are quite variable, median longitudinal extent reached the end of the soil-filled channel (−75 mm) for the Bacteria + Protists treatment, but extended just 10 mm down the root for the Bacteria Only treatment. The high variability in longitudinal extent at Week 3 may be due in part to differing morphology of the root systems that naturally give rise to varying opportunities for bacterial colonization.

**Figure 4.**
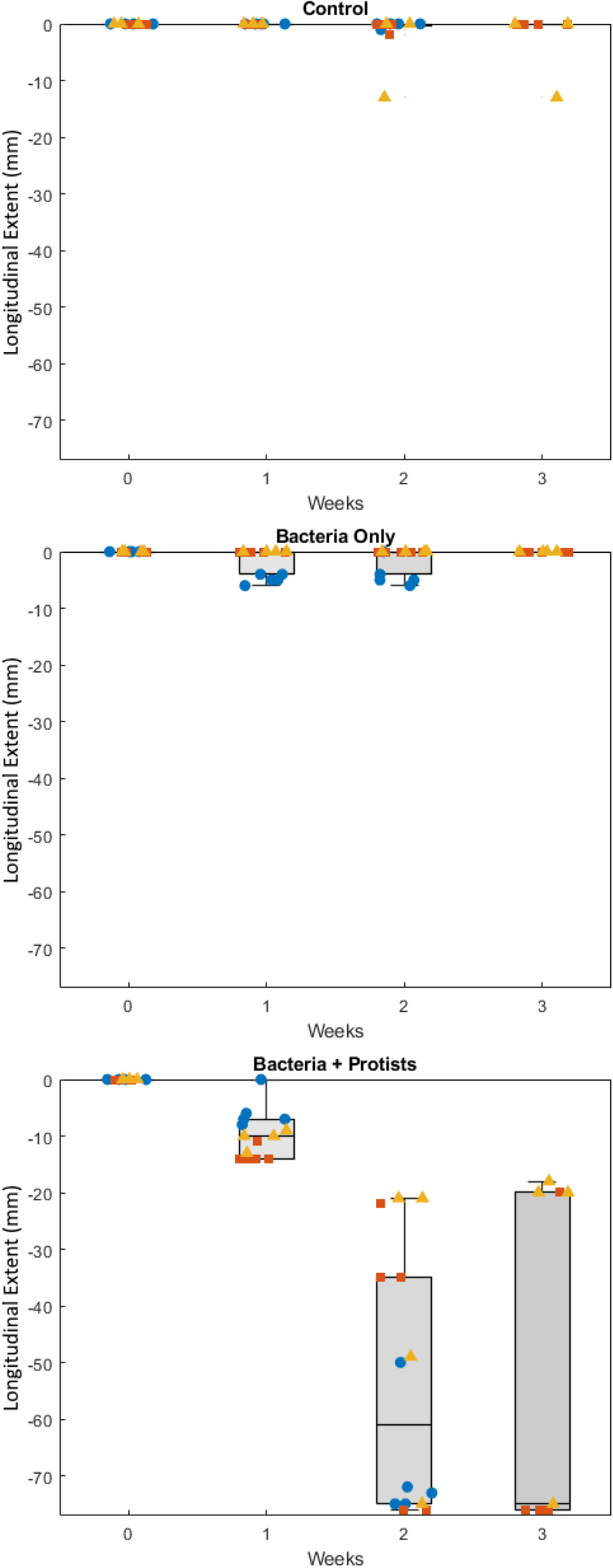
Longitudinal extent (the greatest depth where fluorescent pixel fraction > 10%) by treatment and time. The longitudinal direction is along the major axis of root growth.Experimental runs are grouped by color (Run 1 = blue circles: Run 2 = red squares; Run 3 = yellow triangles). Top and bottom edges of each box represent upper and lower quartiles, and the inside line is the sample median. Whiskers represent the non-outlier minimum and maximum values. Outliers were determined as values more than 1.5 times the inner quartile range beyond the box limits. Treatments are: Control (C) n=13; Bacteria Only (BO) n=14; Bacteria + Protists (BP) n=14.

Roots branch as they grow and infectible root hairs can be found near the tips of all growing roots; therefore, bacterial displacement *across* the major axis of root growth is also important to promote performance of bacterial soil amendments. A plot of lateral extent (the greatest value of fluorescent pixel fraction at any depth, converted to an equivalent width) by replicate and time shows the effect of protists on spreading bacteria across the root system (Figure 5). For the Bacteria + Protists treatment, the median lateral extent was about 8 mm by Week 2, and about 12 mm by Week 3. These lateral extents are substantially greater than those observed in the Bacteria Only treatment, where the median value of lateral extent was about 1 mm, and did not change much over time.

**Figure 5.**
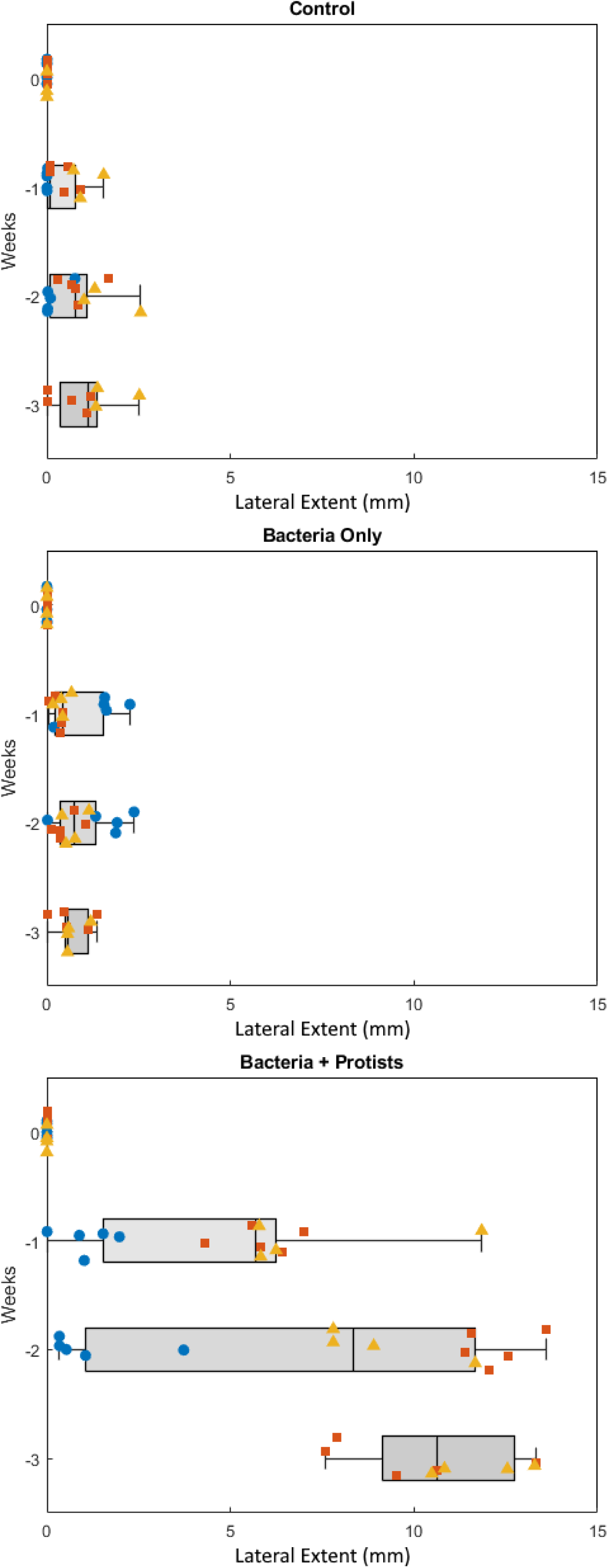
Lateral extent (the largest fluorescent pixel fraction at any depth converted to an equivalent width) of the fluorescence signal by treatment and time. The lateral direction is across the major axis of root growth. Experimental runs are grouped by color (Run 1 = blue circles: Run 2 = red squares; Run 3 = yellow triangles). Top and bottom edges of each box represent upper and lower quartiles, and the inside line is the sample median. Whiskers represent the non-outlier minimum and maximum values. Outliers were determined as values more than 1.5 times the inner quartile range beyond the box limits. Treatments are: Control (C) n=13; Bacteria Only (BO) n=14; Bacteria + Protists (BP) n=14.

Even greater longitudinal and lateral extents may have been observed in a different plant growth chamber, but as the restrictions of our setup constrain all treatments equally, the platform clearly shows the effect of protists on fluorescence signal displacement. Statistical analysis of these results shows that the differences in longitudinal extent and lateral extent among the different treatments are highly statistically significant. Binary unpaired t-tests were done for all combinations of treatment (Bacteria Only, Bacteria + Protists, and Control) for all points in time. Both vertical and horizontal distances are significantly higher in the Bacteria + Protists treatment than in the Bacteria Only treatment or in the Control in every week (Bonferroni corrected p < 0.001) (Table 2). Meanwhile, no significant differences are found between the BO treatment and Control in any week (Bonferroni corrected p > 0.1).

**Table 2:** Pairwise significance (p-values^*^) comparing longitudinal extent (i.e., with plant depth, see Figure 4) and lateral extent (see Figure 5) by treatment and by week. Treatment abbreviations are Control (C), where no microbes are added; Bacteria Only (BO); and Bacteria + Protists (BP).

|  | Treatment Pairs | Week 1 | Week 2 | Week 3 |
| --- | --- | --- | --- | --- |
| Longitudinal Extent |  |  |  |  |
|  | BO v. C | p=0.120 | p=0.869 | p=0.351 |
|  | BP v. C | p<0.001*** | p<0.001*** | p<0.001*** |
|  | BP v. BO | p=0.001*** | P<0.001*** | p<0.001*** |
| Lateral Extent |  |  |  |  |
|  | BO v. C | p=0.138 | p=0.590 | p=0.349 |
|  | BP v. C | p<0.001*** | p<0.001*** | p<0.001*** |
|  | BP v. BO | p=0.001*** | p<0.001*** | p<0.001*** |

p-values of unpaired t-tests, two-sided, unequal variance with Bonferroni correction for multiple hypothesis tests
\*\*\* p<0.01, \*\* p<0.05, \* p<0.10

These comparisons above were performed by grouping all experiments together.However, we detailed important differences in how the bacteria were added, whether the protists were “awake” when added, and in the watering condition—these methodological variations certainly impacted bacterial transport. To test for the effect of experimental conditions (i.e., “Experimental Run”) on our results, we performed a linear regression and found that even when controlling for experimental run, the treatment effect of protists on longitudinal and lateral extent is still highly significant (Table S1).

Although despite methodological differences protists always promote bacterial dispersal, we observed interesting differences in the displacement patterns with experimental run that are worth considering. In Experimental Run 1, where soil conditions were dry, the fluorescence signal was found primarily along the roots. In Experimental Runs 2 and 3 we observed a more prominent, wider, and somewhat shallower pattern of fluorescence. In examining the results quantitatively, the most significant difference is the lateral spread of the fluorescence signal. For Experimental Run 1, median lateral extents were 1.1 mm (Week 1) and 1.2 mm (Week 2). For Runs 2 and 3, where plants were watered nearly twice as often, median lateral extent were 5.8 mm (Week 1) and 12.2 mm (Week 2) and 7.4 mm (Week 1) and 9.0 mm (Week 2), respectively. These differences are statistically significant (p=0.02 or less). In terms of longitudinal extent, differences apparent at Week 1 were not significant by Week 2 (Table S2). In Experimental Run 1, the reduced water application rate may have been associated with a less-branched root morphology, as the plant sent roots downward in search of water, as well as fewer contiguous water-saturated pathways through the soil pore volume other than directly along the root itself. Both of these factors would tend to focus microbial activity immediately around the roots and its associated mucilage and promote longitudinal transport at the expense of lateral.

### Protists enhance dispersal of viable bacteria along root systems

After imaging was complete, µ-rhizoslides were disassembled and the root systems (root tissue plus adhering soil) were divided into shallow and deep segments. The bacterial abundance in the shallow segment, as determined by plating for CFUs, averaged 1.02 × 10^7^ in the Bacteria Only treatment and 1.44 × 10^7^ in the Bacteria + Protists treatment (Table 3, Figure 6). These abundances are not statistically significant (p = 0.315). In other words, despite predation by protists, there is a similar total bacterial population in the shallow root system of the Bacteria Only and Bacteria + Protists treatments, implying that access to habitat niches and/or substrates may limit bacterial population size in the shallow portion of our setup and not the effects of predation.

**Table 3:** Bacterial abundance (×10^7^ as colony forming units, or CFUs) averaged over all runs and replicates by root system segment (i.e., shallow v. deep) and treatment with statistical significance (p-value). Treatment abbreviations are Control (C), where no microbes are added; Bacteria Only (BO); and Bacteria + Protists (BP).

| Treatment | Shallow | Deep | Total | Shallow v. Deep<sup>+</sup> |
| --- | --- | --- | --- | --- |
| C | 0 | 0 | 0 | --- |
| BO | 1.02 | 0 | 1.02 | p=0.001*** |
| BP | 1.44 | 0.458 | 1.90 | p=0.008** |
| BP v. BO<sup>*</sup> | p=0.315 | p=0.013** | p=0.080* |  |

<sup>\*</sup>BP v. BO: p-values of unpaired t-tests, two-sided, unequal variance
<sup>+</sup>Shallow v. Deep: p-values of paired t-tests, two-sided
\*\*\* $p < 0.01$ , \*\* $p < 0.05$ , \* $p < 0.10$

**Figure 6.**
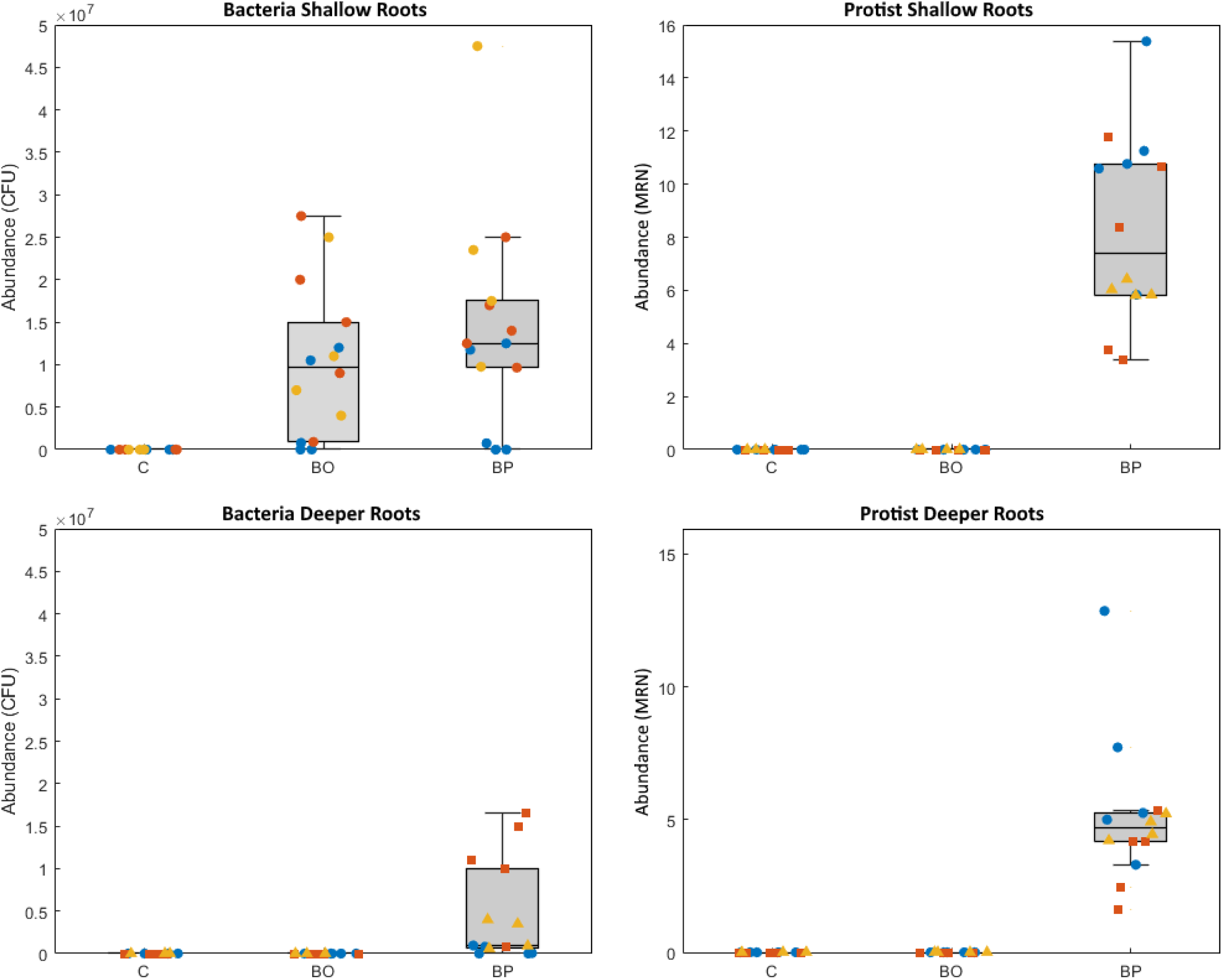
Bacteria and protist abundance in the top (“shallow roots”) and bottom (“deeper roots”) segments of *M. truncatula* root systems. Bacteria abundance is represented by average colony forming units (CFUs) in plated serial dilutions of the root system extract. Protist abundance is represented by average minimum recovered number (MRN) of protists in propagated serial dilutions of the root system extract. Experimental runs are grouped by color (Run 1 = blue circles: Run 2 = red squares; Run 3 = yellow triangles). Top and bottom edges of each box represent upper and lower quartiles, and the inside line is the sample median. Whiskers represent the non-outlier minimum and maximum values. Outliers were determined as values more than 1.5 times the inner quartile range beyond the box limits. Treatments are: Control (C) n=13; Bacteria Only (BO) n=14; Bacteria + Protists (BP) n=14.

In the deeper root system, we found 0.458 × 10^7^ CFU bacteria in the Bacteria + Protists treatment (Table 3, Figure 6). versus 0 CFU recovered in the Bacteria Only treatment. We only recovered bacteria from deeper roots in replicates where protists were present. We found here that bacteria were unable to reach the deeper roots in our setup without the assistance of protists. These results support the fluorescence data in showing that protists not only enhance displacement of a bacteria-associated fluorescence signal but also seem to directly enhance the spatial distribution of recoverable, viable bacteria in growing root microcosms.

For completeness, we also enumerated protists in root system segments. We only recovered protists in the Bacteria + Protists treatment, and report comparable numbers from both shallow and deep root system segments (Table 4 and Figure 6). This result implies that protists fully disperse within the soil system by the end of the experiment.

**Table 4:** Protist abundance averaged across all runs and replicates. Abundance is estimated as minimum recovered number (MRN) of propagated extract dilutions by root system segment (i.e., shallow v. deep) and treatment with statistical significance (p-value).

| Treatment | Shallow | Deep | Total | Shallow v. Deep<sup>+</sup> |
| --- | --- | --- | --- | --- |
| C | 0 | 0 | 0 | --- |
| BO | 0 | 0 | 0 | --- |
| BP | 233 | 185 | 419 | p=0.026** |

<sup>+</sup>Shallow v. Deep: p-values of paired t-tests, two-sided
\*\*\* $p < 0.01$ , \*\* $p < 0.05$ , \* $p < 0.10$

### Conclusions, Limitations, and Applications

While previous work has shown the potential for using protists as vehicles for transporting particles (20), here we demonstrate the potential for protists to transport viable beneficial bacteria in a system containing a living plant and real soil. In this report, we show protist-facilitated transport increases abundance of viable bacteria in the deeper root system segment despite losses due to grazing. The imaging experiments showed distinct patterns of fluorescence displacement both longitudinally and laterally over time – but only when protists were present. In the absence of protists, there was no statistically significant change in the fluorescence signal with depth or breadth versus the control, and the imaging showed that the fluorescence signal remained within 5 mm of the inoculation site for the duration of the experiment.

Because fluorescent protein levels will likely vary between bacterial cells due to differences in plasmid number, growth rate, and cell size, the fluorescence signal is most conservatively interpreted as an indicator of bacteria position, and not a direct measurement of bacterial abundance or biomass. In addition, our fluorescence results may have been limited because the fluorescent plasmid pCMB35, which is usually maintained with 5 μg/mL tetracycline, may have not been retained because tetracycline was not added to the soil in the µ-rhizoslides. We opted not to add tetracycline because of the possibility that it could adversely impact plant health and development. Initially, we attempted to use a different fluorescent protein-expressing *S. meliloti* strain with a plasmid that is maintained without selection, L5-30/pDG77, but the resulting DsRed fluorescence signal was not compatible with our imaging setup.

For all these reasons, our quantitative approach focused on the extent of the spatial domain that contained an appreciable fluorescent signal (i.e., fluorescent pixel fraction > 0.1 by section). We did not attempt to aggregate total fluorescence as a proxy for abundance or biomass. The choice to emphasize spatial extent of fluorescence and not amount of fluorescence helps to explain why longitudinal and lateral extent are dramatically different between treatments yet we recover similar numbers of bacteria in the top (shallow root system) segment of the Bacteria Only and Bacteria + Protists treatments. In the former, the entire population of bacteria is limited to the initial inoculation site, whereas bacteria in the Bacteria + Protists treatment are spread out over a larger area. Importantly, *S. meliloti* strain Rm1021/pCMB35 was not recovered in the deeper system segment at all unless protists were present. The bacteria appear not to be able to reach that area of the soil channel after three weeks of plant development and growth without the assistance of protists.

The quantities of bacteria and protists recovered in our study represent minimum values for several reasons. First, only bacteria and protists directly associated with roots and root-adhering soil were quantified. The bulk soil, which was not evaluated by these methods, likely contained additional organisms. Also, our methods may tend to under-represent the true microbial abundance, especially for protists. For our MRN assay the true value may be approximately three times higher because of the assumption that the last well of a dilution series initially contained only one protist when in fact it may have contained two or more protists.

However, despite these important methodological details, the trend in our data strongly support the hypothesis that protists enhance dispersal of bacteria along root systems, including bacteria that are recoverable and viable. The effect of these bacteria on the roots and plant health is the subject of a further study.

The protist used in this study, *Colpoda sp*., a ciliate isolated from a legume rhizosphere, is particularly successful in transporting bacteria. Ciliates most often filter feed, a mechanism of promiscuous consumption of food particles and prey. Bacteria consumed by the ciliates are packaged into a phagosome which is processed and eventually egested from the protist; a process that is not well characterized. It is thought that egested phagosomes break down in the soil and these may release viable bacteria which could go on to colonize the area where they were released. How often bacteria can survive intake and egestion, and whether some species are adept at this in not clearly understood. Some bacterial species, for example *Campylobacter jejuni, Listeria spp*. and *Mycobacterium* spp. are known to escape digestion and colonize the cytoplasm of protists that consumed them (38, 49). Ciliates have also been shown to emerge from feeding on bacterial biofilms with a matrix of bacteria and EPS stuck to their cell bodies (20). When the protists travel along plant roots consuming bacteria, both egestion of viable bacteria and physical translocation of bacteria may contribute to protist-assisted transport.

The technology described here provides a simple, reproducible experimental system which has the potential to aid in the understanding of rhizosphere biology. Of particular interest is bacterial movement in the context of screening a variety of different bacteria/protist/plant/soil combinations for their effects on such movement. For example, protist transportation of *Sinorhizobium sp*. and its effect on nodulation can be studied using this system and parameters that affect transport such as water saturation, soil composition and soil porosity can be readily altered then tested and quantitated. Protist-facilitated transport may also be adapted for targeted delivery of specialized microbes or encapsulated agrochemicals. With increased understanding of protist-facilitated transport mechanisms and limitations under reproducible yet realistic conditions we can begin developing new methods to support sustainable agriculture and have increasing confidence of their utility for use in the field.

## Acknowledgements

This work was supported by DOE award DE-SC0014522 (to LMS and DJG), NSF award 1605816 (to LMS and DJG), and USDA National Institute of Food and Agriculture award 2016-67013-24412 (to DJG and LMS).

