## Supplementary material for "Soil protists can actively redistribute beneficial bacteria along *Medicago truncatula* roots": Supp Tables 1 and 2, Supp Figures 1 and 2

**Table S1:** Determinants of longitudinal and lateral extent by week (linear regression estimates). Control Treatment and Experimental Run 1 are the omitted variables. Treatment abbreviations: Bacteria Only (BO); Bacteria + Protists (BP).

| | Longitudinal Extent | | | Lateral Extent | | |
|---|---|---|---|---|---|---|
| | Week 1 | Week 2 | Week 3 | Week 1 | Week 2 | Week 3 |
| Treatment BO | -1.740 | -0.513 | 0.548 | 0.270 | 0.098 | -0.389 |
| | (1.074) | (5.147) | (8.350) | (0.679) | (1.016) | (0.613) |
| Treatment BP | -9.803*** | -52.963*** | -50.177*** | 4.094*** | 6.544*** | 9.582*** |
| | (1.074) | (5.147) | (8.350) | (0.679) | (1.016) | (0.613) |
| Experimental Run 2 | -0.999 | 7.996 | -10.095 | 1.364** | 3.578*** | -1.050** |
| | (1.017) | (4.873) | (6.825) | (0.643) | (0.962) | (0.501) |
| Experimental Run 3 | -0.001 | 9.744* | -- | 2.168*** | 2.827** | -- |
| | (1.107) | (5.305) | | (0.700) | (1.047) | |
| Constant | 0.385 | -6.553 | 5.061 | -0.616 | -1.254 | 1.681*** |
| | (0.952) | (4.563) | (7.416) | (0.602) | (0.900) | (0.545) |
| N | 41 | 41 | 26 | 41 | 41 | 26 |
| $R^2$ | 0.731 | 0.803 | 0.705 | 0.617 | 0.666 | 0.943 |
| F | 24.423 | 36.775 | 17.487 | 14.489 | 17.907 | 120.682 |

*Standard errors are in parentheses*
**** p<0.01, ** p<0.05, * p<0.10*

**Table S2:** Mean values (standard deviation in parenthesis) of longitudinal and lateral extent by week and experimental run with significance of pairwise t-tests. Treatment abbreviations are Bacteria + Protists (BP) and Bacteria Only (BO).

| | Longitudinal Extent | | Lateral Extent | |
|---|---|---|---|---|
| | Week1 | Week 2 | Week 1 | Week 2 |
| **Treatment BP** | | | | |
| Experimental Run 1 | -5.594 | -68.941 | 1.072 | 1.197 |
| | (3.206) | (10.696) | (0.746) | (1.444) |
| Experimental Run 2 | -13.386 | -48.749 | 5.826 | 12.222 |
| | (1.340) | (25.364) | (1.015) | (0.894) |
| Experimental Run 3 | -10.489 | -41.456 | 7.423 | 9.043 |
| | (1.730) | (25.915) | (2.954) | (1.825) |
| t-tests | | | | |
| Exp 2 v. 3 | p=0.036** | p=0.686 | p=0.366 | p=0.032** |
| Exp 1 v. 3 | p=0.025** | p=0.121 | p=0.020** | p=0.001*** |
| Exp 1 v. 2 | p=0.003*** | p=0.158 | p<0.001*** | p<0.001*** |
| **Treatment BO** | | | | |
| Experimental Run 1 | -4.795 | -3.996 | 1.440 | 1.501 |
| | (0.836) | (2.343) | (0.759) | (0.910) |
| Experimental Run 2 | 0 | 0 | 0.360 | 0.535 |
| | (0) | (0) | (0.083) | (0.360) |
| Experimental Run 3 | 0 | 0 | 0.412 | 0.709 |
| | (0) | (0) | (0.208) | (0.328) |
| t-tests | | | | |
| Exp 2 v. 3 | | | p=0.659 | p=0.475 |
| Exp 1 v. 3 | p<0.001*** | p=0.019** | p=0.036** | p=0.129 |
| Exp 1 v. 2 | p<0.001*** | p=0.019** | p=0.033** | p=0.076* |

*p-values of unpaired t-tests, two-sided, unequal variance*

**** p<0.01, ** p<0.05, * p<0.10 with Bonferroni correction for 12 multiple hypothesis tests*

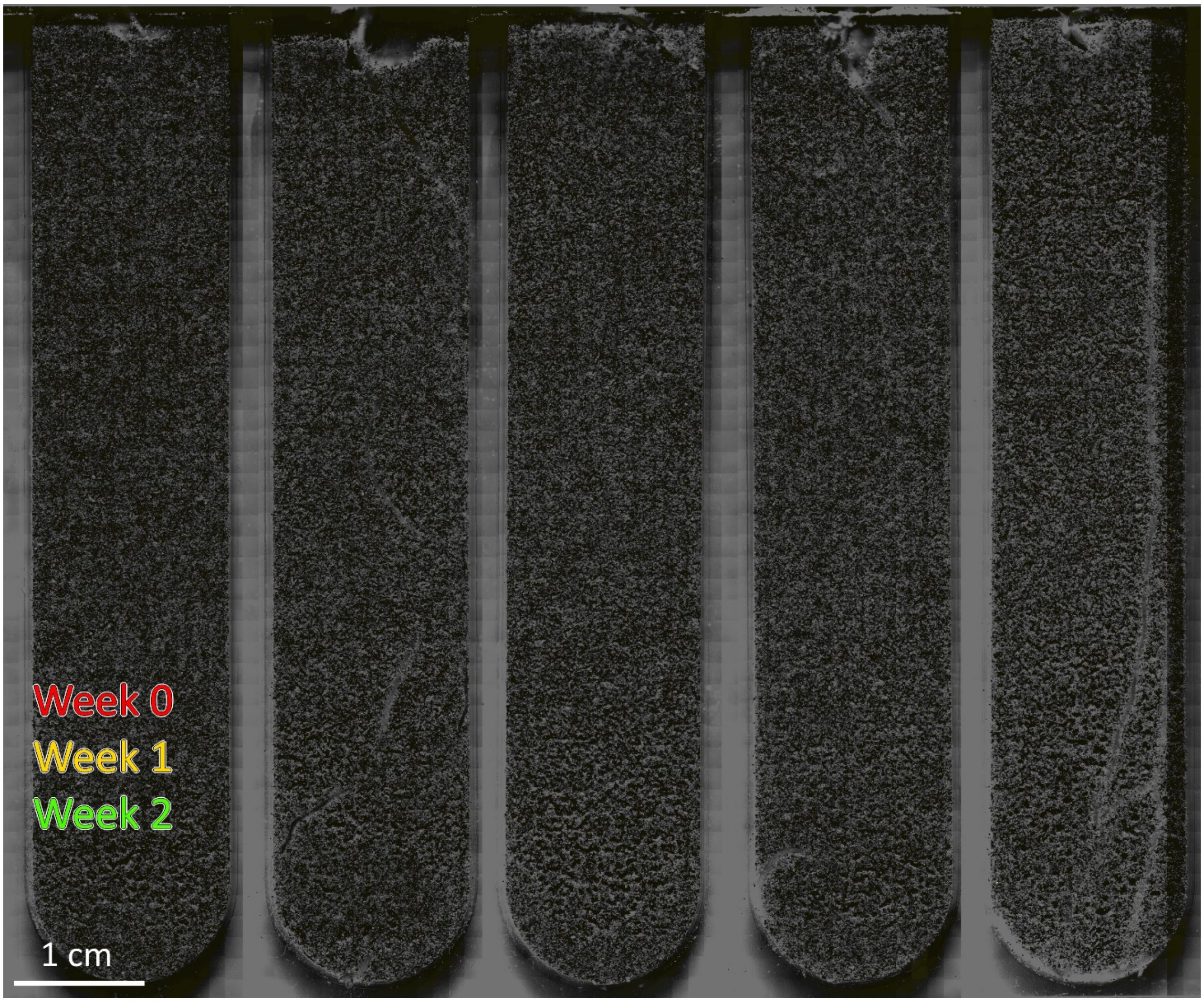


**Figure S1.1A** Composite brightfield (at last imaging date) and fluorescence (false-colored by week as indicated) images of soil channels from **Experimental Run 1: Control replicates**. In this experiment, five soil and plant replicates were inoculated with 10 μL buffer: no microbes were added.

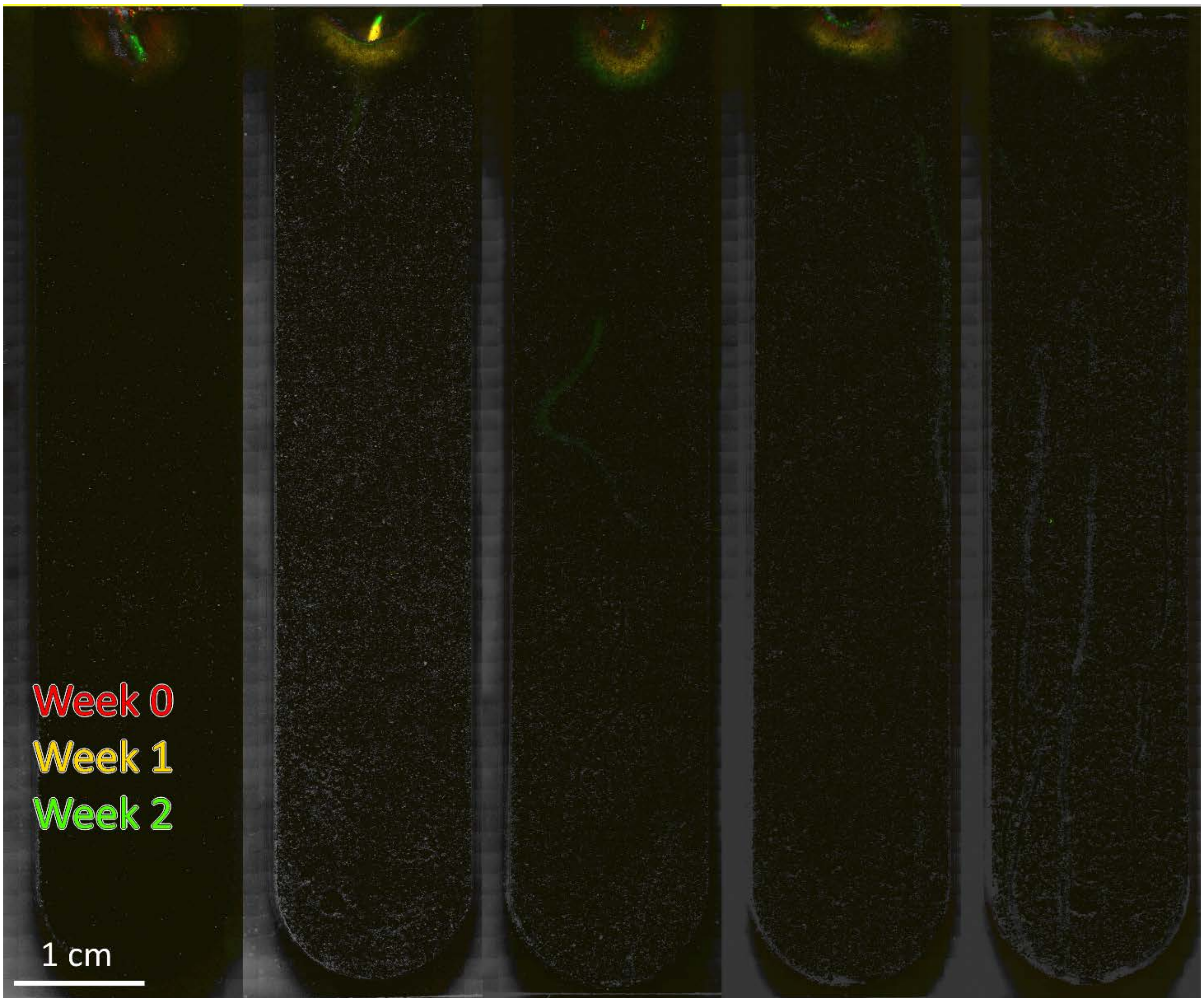


**Figure S1.1B** Composite brightfield (at last imaging date) and fluorescence (false-colored by week as indicated) images of soil channels from **Experimental Run 1: Bacteria Only replicates**. In this experiment, five soil and plant replicates were inoculated with 10 µL buffer containing $10^7$ CFU bacteria.

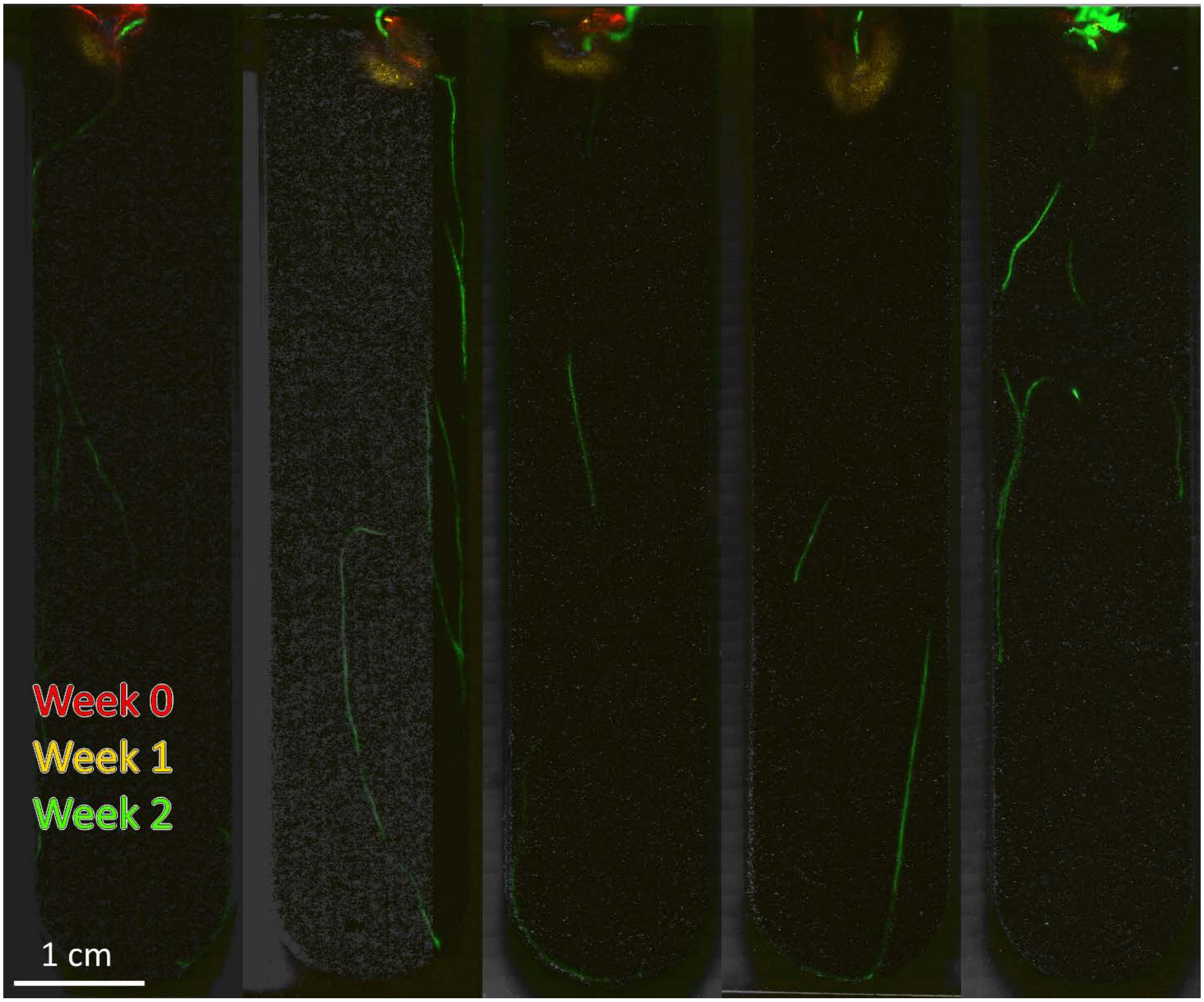


**Figure S1.1C** Composite brightfield (at last imaging date) and fluorescence (false-colored by week as indicated) images of soil channels from **Experimental Run 1: Bacteria + Protists replicates**. In this experiment, five soil and plant replicates were inoculated with 10 µL buffer containing $10^7$ CFU bacteria plus $1.5 \times 10^3$ *Colpoda sp*. protists.

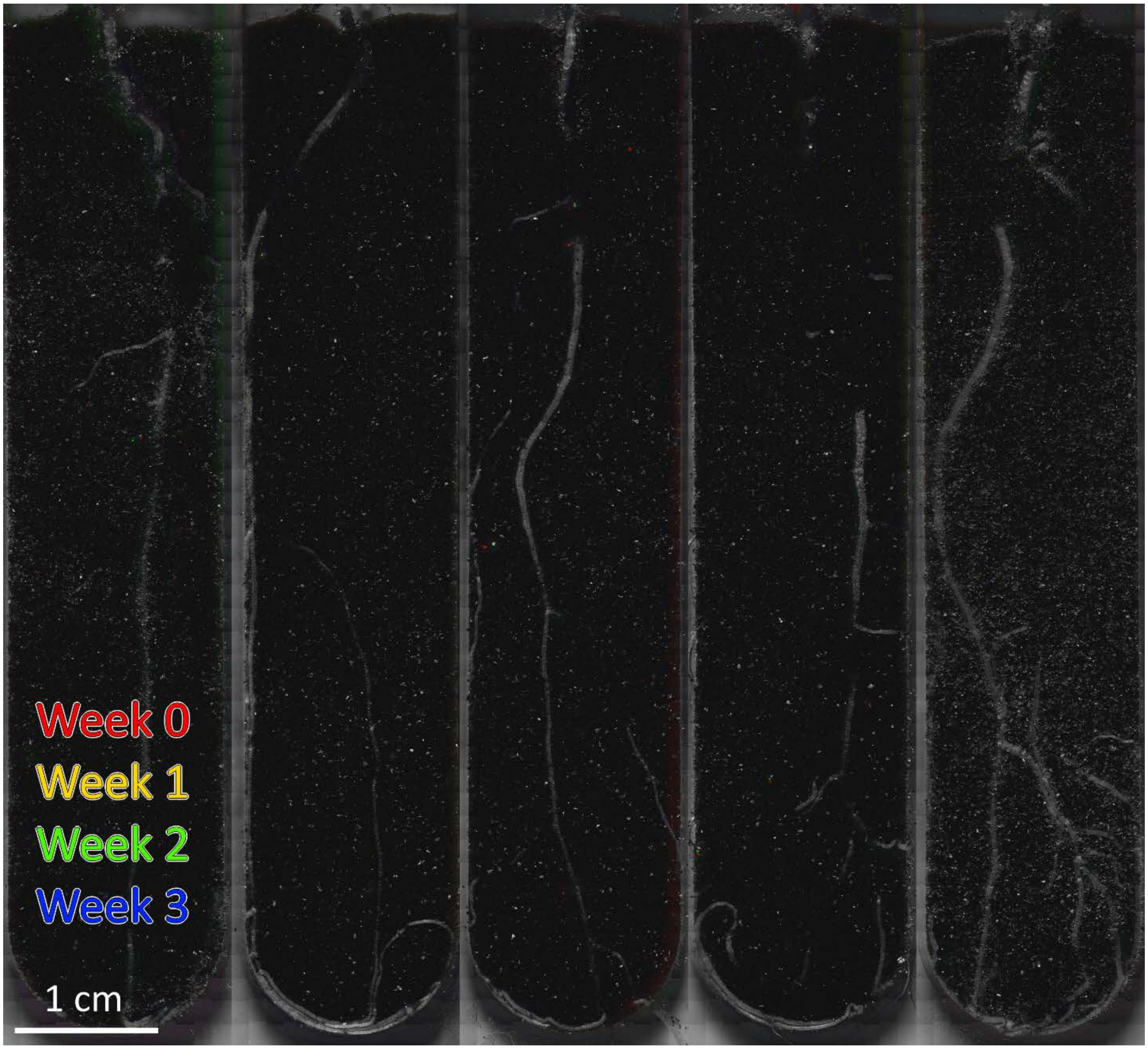


**Figure S1.2A** Composite brightfield (at last imaging date) and fluorescence (false-colored by week as indicated) images of soil channels from **Experimental Run 2: Control replicates**. In this experiment, five soil and plant replicates were inoculated with 10 μL buffer: no microbes were added.

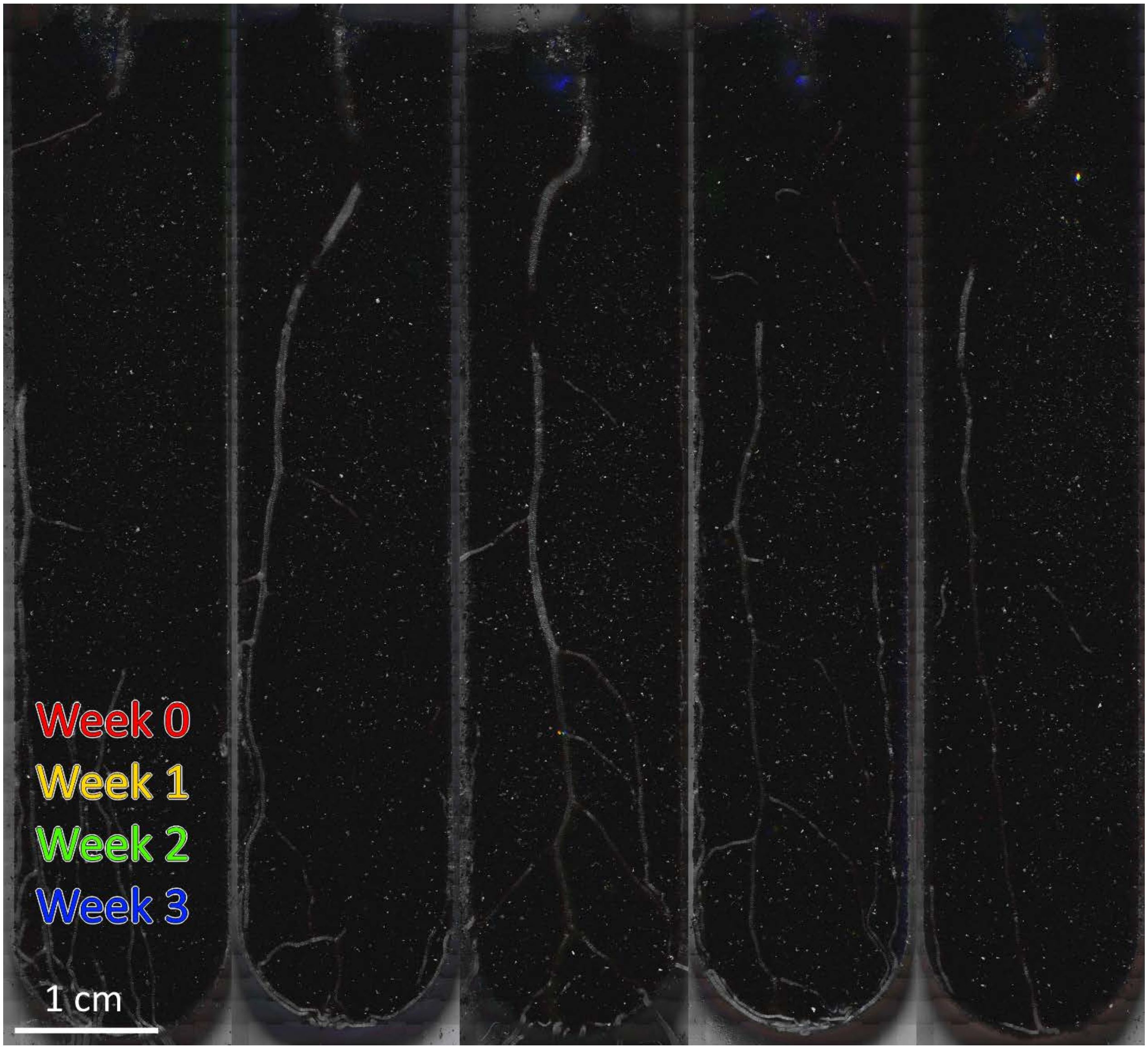


**Figure S1.2B** Composite brightfield (at last imaging date) and fluorescence (false-colored by week as indicated) images of soil channels from **Experimental Run 2: Bacteria Only replicates**. In this experiment, five soil and plant replicates were inoculated with 10 µL buffer containing $10^7$ CFU bacteria.

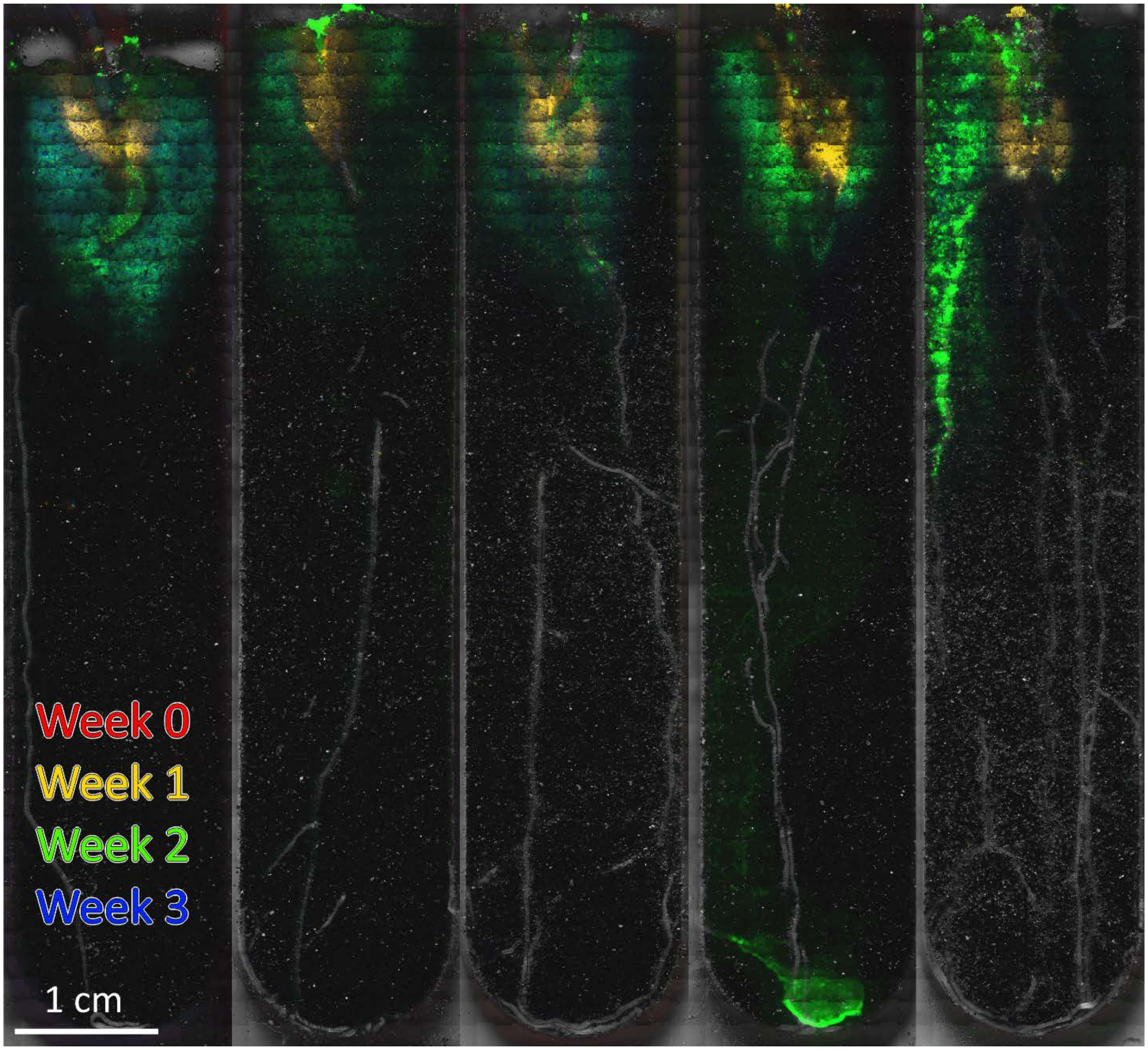


**Figure S1.2C** Composite brightfield (at last imaging date) and fluorescence (false-colored by week as indicated) images of soil channels from **Experimental Run 2: Bacteria + Protists replicates**. In this experiment, five soil and plant replicates were inoculated with 10 µL buffer containing $10^7$ CFU bacteria plus $1.5 \times 10^3$ *Colpoda sp*. protists.

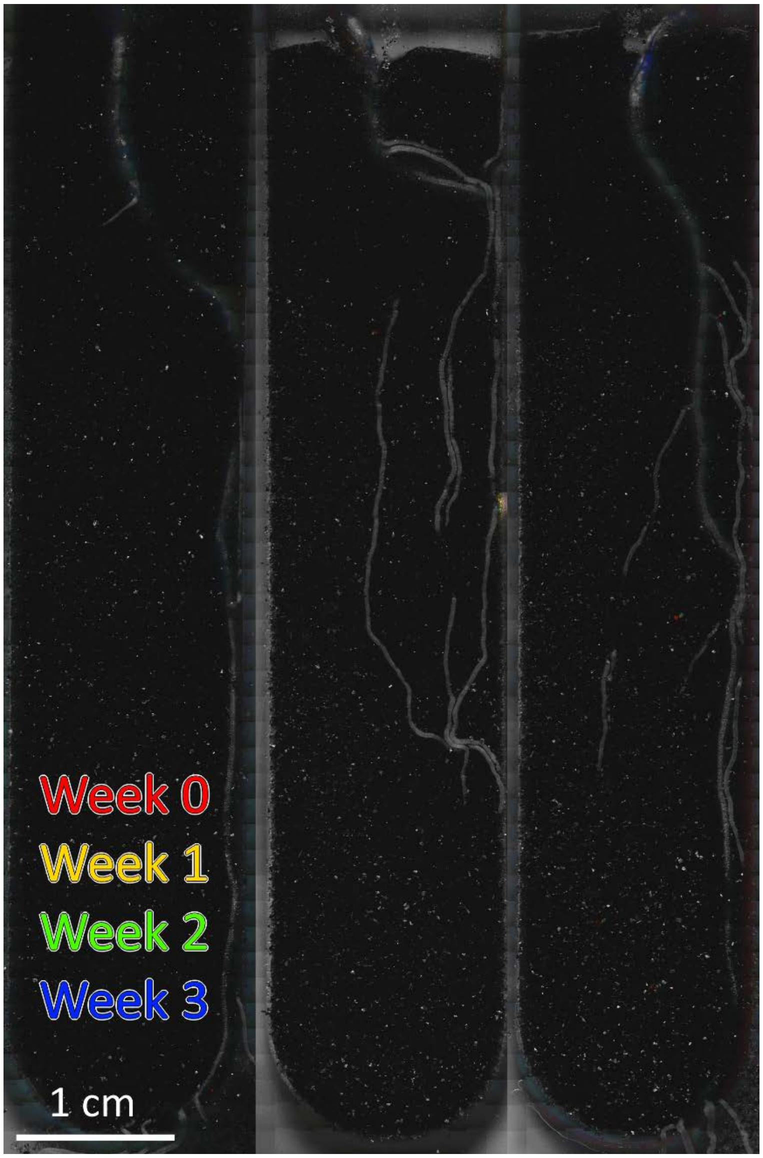


**Figure S1.3A** Composite brightfield (at last imaging date) and fluorescence (false-colored by week as indicated) images of soil channels from **Experimental Run 3: Control replicates**. In this experiment, three soil and plant replicates were inoculated with 10 μL buffer: no microbes were added.

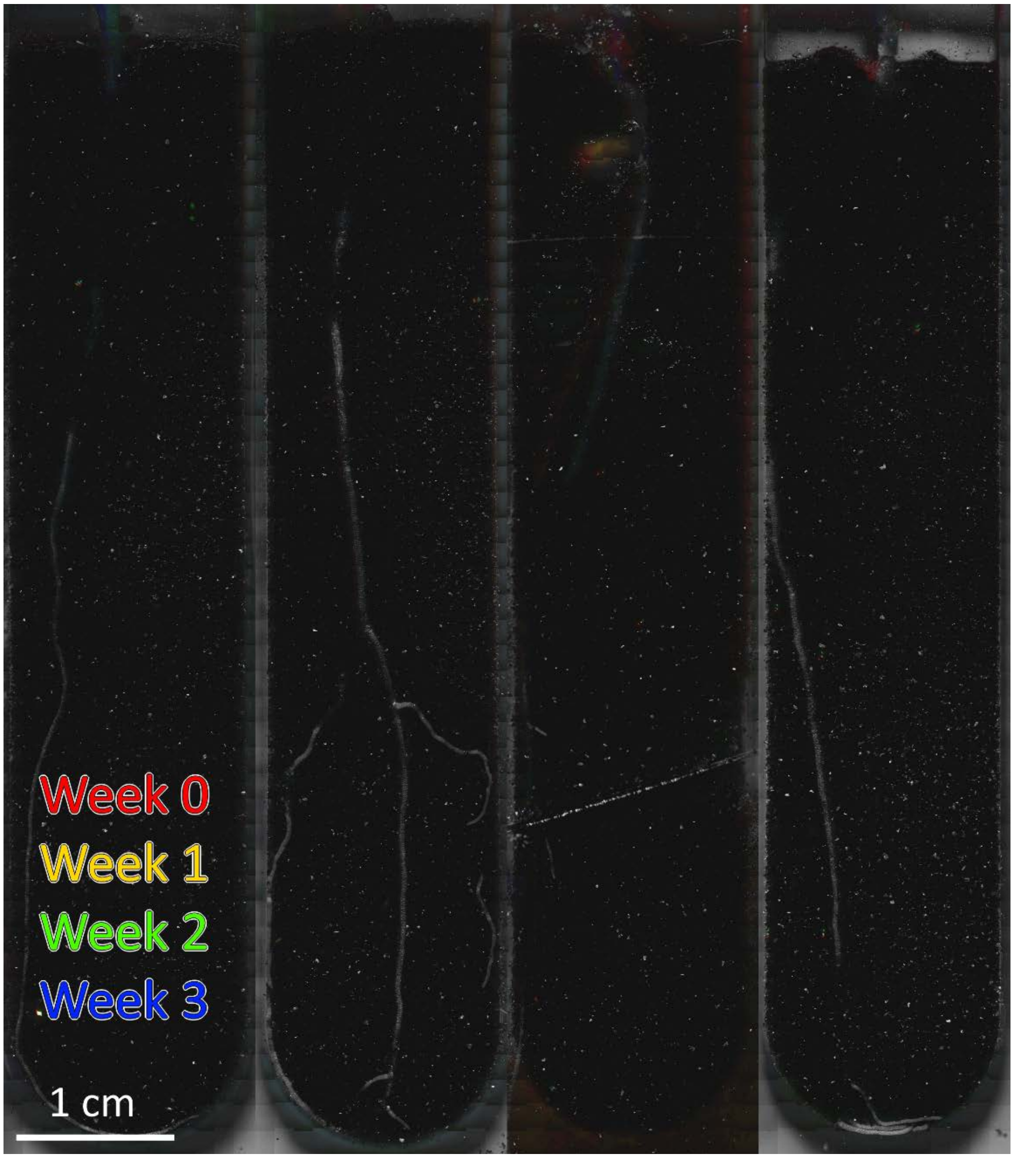


**Figure S1.3B** Composite brightfield (at last imaging date) and fluorescence (false-colored by week as indicated) images of soil channels from **Experimental Run 3: Bacteria Only replicates**. In this experiment, four soil and plant replicates were inoculated with 10 µL buffer containing $10^7$ CFU bacteria.

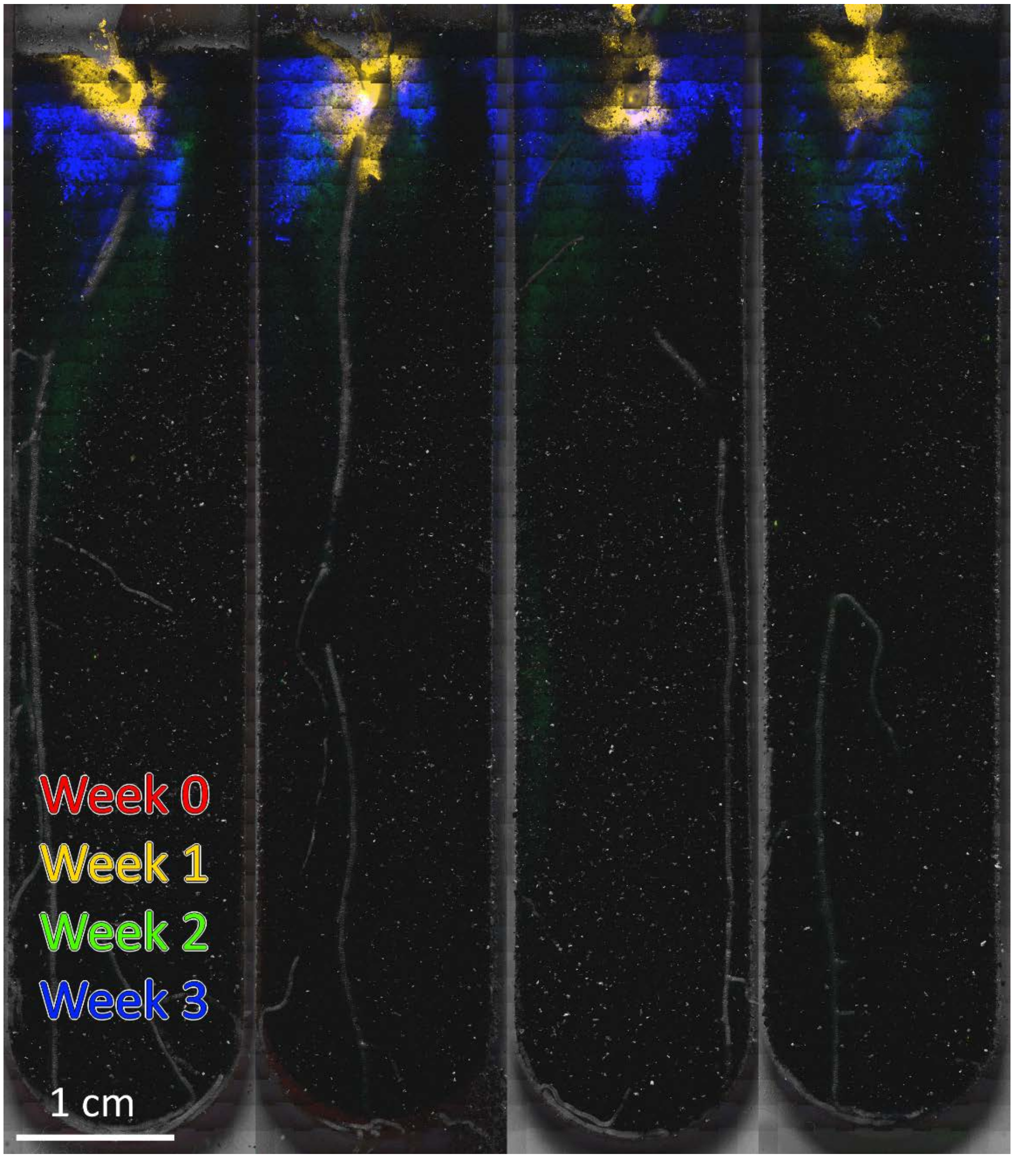


**Figure S1.3C** Composite brightfield (at last imaging date) and fluorescence (false-colored by week as indicated) images of soil channels from **Experimental Run 3: Bacteria + Protists replicates**. In this experiment, four soil and plant replicates were inoculated with 10 μL buffer containing $10^7$ CFU bacteria plus $1.5 \times 10^3$ *Colpoda sp*. protists.

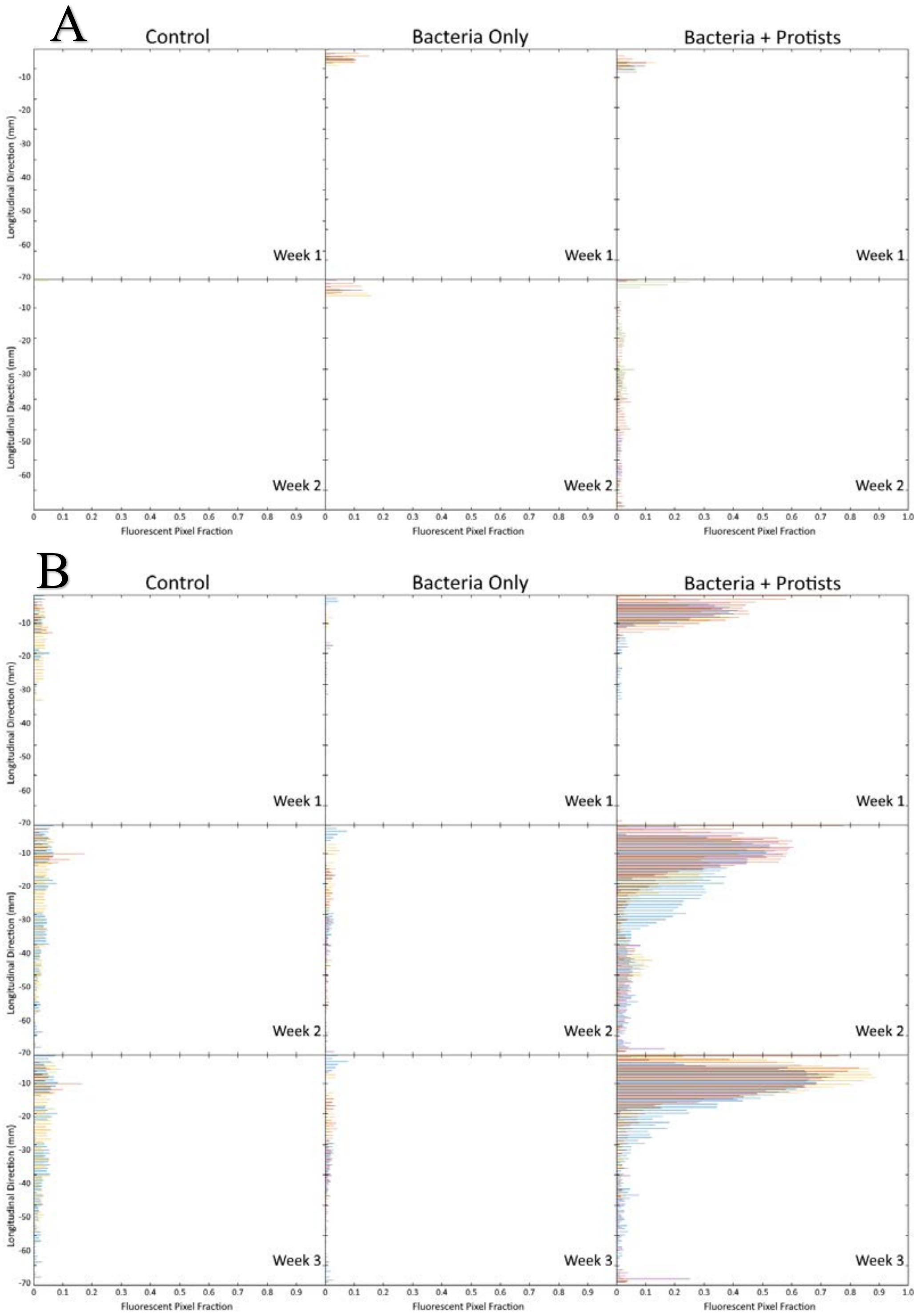


**Figure S2 Spatial distribution of fluorescence signal over time.** For each treatment (arranged above in columns), the entire soil channel was imaged at different points in time (Week 0, Week 1, Week 2, Week 3). Here, "Fluorescent Pixel Fraction" (the ratio of non-zero pixels to total pixels) was computed for binned strips (each measuring 1 mm high and 15 mm wide) of the soil channel after background was subtracted (Week 0) and plotted with distance in the longitudinal direction (i.e., along the axis of the growing root). Replicates distinguished by color and panels correspond with: A) Experimental Run 1; and B) from Experimental Run 3.
